# Drebrin forms a cortical hub that connects actin, microtubules, and clathrin for endocytic transport of β2 integrin

**DOI:** 10.64898/2026.03.12.711275

**Authors:** Pasquale Cervero, Sven Hey, Kathrin Weber, Bryan Barcelona, Robert Herzog, Stefan Linder

## Abstract

Using primary human macrophages, we analyze the molecular machinery that enables direct contact of microtubule +tips with podosomes, and its connection to clathrin-based endocytosis. We show that the podosome cap protein drebrin, together with its +tip-localized binding partner EB3, regulates podosome-microtubule contact. Importantly, drebrin depletion leads to reduced endocytosis, particularly of β2 integrin at podosomes. This is based on drebrińs interaction with β2 integrin, and its direct binding of clathrin heavy chain via a canonical clathrin box and a novel box, which we identify also in other mammalian proteins. Our data enable a model of the molecular machinery that regulates direct contact between podosomes and microtubule +tips. Moreover, we provide evidence for spatial selectivity in endocytosis of β integrins, while also revealing a more general principle of endocytic trafficking regulation, through drebrin working as a multivalent, actin-based hub that links the clathrin system with microtubule-based transport.

## Introduction

Microtubules (MTs) constitute the cellular long-range transport system that enables trafficking of cargo to and from the cell cortex, thereby regulating exo- and endocytosis ^1^. Moreover, contact with microtubules influences the formation and dynamics of integrin-based adhesion at the cell cortex such as focal adhesions and podosomes ^2^. However, the molecular mechanisms coupling microtubule targeting of adhesions to exo-/endocytic events remain poorly understood.

Podosomes are cortical adhesion and invasion structures in several cell types, notably macrophages ^3^, immature dendritic cells ^4^, osteoclasts ^5^, endothelial cells ^6^ or neural crest cells ^7^,. They mediate extracellular matrix-cytoskeleton linkage through receptors such as integrins^8^ or CD44 ^9^, but are also active in matrix degradation ^10^, as well as sensing of substrate properties such as rigidity ^11,12^ and topography ^13^. Together with invadopodia of cancer cells, they form the subgroup of invadosome adhesions ^14^.

A hallmark of podosomes is their F-actin-rich core, based on Arp2/3 activity ^3,15^, which is surrounded by a discontinuous ring ^16^ of plaque proteins such as talin, vinculin and paxillin ^17–20^. Two sets of unbranched actin filaments have been identified: lateral filaments that connect the top of the core to the ring ^21^, and the dorsal connecting cables that link individual podosomes into higher-ordered groups ^21–23^.

A further substructure, the podosome cap, positioned atop of the actin core, has emerged as a regulatory module that acts on key podosome functions such as oscillation or matrix degradation ^24–26^. The cap represents the lateral unbranched actin filaments, together with their binding partners, which are mostly proteins involved in actin crosslinking or bundling ^25^. Importantly, the cap has been proposed to play a critical role in regulating the contact between podosomes and microtubules, serving as a hub for incoming vesicles ^27^, and as a mechanoregulatory module that enables the mechanostimulation or recruitment of proteins at the podosome ring ^28^. Moreover, both functions may not be mutually exclusive ^29^.

MTs impact on numerous aspects of podosome dynamics and functions such as formation ^30^, dissolution ^31^ and matrix degradation ^10,32^, through the transport of cargo such as matrix metalloproteinases ^10^ or integrins ^33^, by motors like kinesin-1 and-2 or dynein ^10^, KIF1C ^31,34^ or KIF9 ^35^. The capture of MT +tips at podosomes is therefore crucial for the delivery and recycling of regulatory and effector proteins.

Previous findings revealed an indirect mechanoregulation of podosomes by microtubules, through KANK proteins and the RohGEF GEFH1 ^2^. Here, we screened mostly for regulators of direct MT-podosome contact. The cap structure surrounds the podosome core and is exposed to the cytoplasm ^25^, rendering cap proteins such as LSP1 ^24^, INF2 ^26^, supervillin ^22^, or the here newly identified cap protein drebrin, compelling candidates for such an investigation. Our screen also included components of cortical docking complexes (CDCs) that consist of cortex-associated proteins such as liprin, ELKS, and LL5β ^36^. CDCs are linked to microtubules via CLASPs ^37^, and to integrin-based adhesions via KANK1 binding to talin^38^. Further candidates included IQGAP1, a linker between MT +tips and actin (Watanabe et al., 2004), found in microglia podosomes ^39^, and PAK4, a modulator of actin and microtubule dynamics, which binds to the Rho-family guanine nucleotide exchange factor (GEF), GEF-H1^40^ and localizes to the ring structure of macrophage podosomes ^41^.

Recent findings suggest a connection between invadosomes and endocytic uptake of cell-surface associated proteins or particles. Examples include podosome-like structures facilitating β3 integrin endocytosis on RGD-containing lipid bilayers ^42^, PI(3,4)P_2_-mediated membrane tubulation at invadopodia enhancing integrin endocytosis ^43^, and macrophage podosomes being hijacked as endocytic entry sites by the human immodeficiency virus (HIV)-1 ^44^. However, the molecular machinery linking invadosomes to endocytosis and subsequent intracellular transport remained undefined. Moreover, the molecular basis for selectivity of β integrin isoforms during clathrin-mediated endocytosis is currently unknown.

We now expand on these findings by identifying the actin-associated protein drebrin as a novel podosome cap component that mediates clathrin-based endocytosis of β2 integrin, which has been shown to be essential for podosome dynamics ^45^. Drebrin contains two actin binding sites, the main one located in the N-terminus, essential for direct binding to F-actin ^46,47^, and an additional cryptic site enabling the bundling of actin filaments, following phosphorylation of the S142 residue by Cdk5 ^48^. Notably, drebrin directly binds to the +tip protein EB3, thus enabling coordinated actin and microtubule dynamics in developing rat embryonic cortical neurons ^48,49^. Importantly, we show that drebrin binds clathrin heavy chain via canonical and non-canonical clathrin boxes in its C-terminus, and that its depletion reduces endocytosis of β2 integrin. These data reveal drebrin as a multivalent hub that links clathrin-dependent endocytosis with microtubule-based transport at cortical actin-rich adhesion sites.

## Results

### Software-based analysis of podosome /+tip contact

In order to analyze contact events between podosomes and microtubule +tips, a TrackMate-based software tool was developed (“ContactAnalyzer”). ContactAnalyzer uses live cell videos, with frame-by-frame detection of both structures and the calculation of circular intersections of their coordinates, with a given, but adjustable, detection diameter (12 px =0.96 µm for podosomes; 5 px =0.4 µm for +tips). For a more detailed description, respective advantages and limitations, see (Fig. 1a,b, Ext. Data Fig. 1), Methods, Discussion, and https://github.com/mbwcode/ContactAnalyzer.

**Figure 1.**
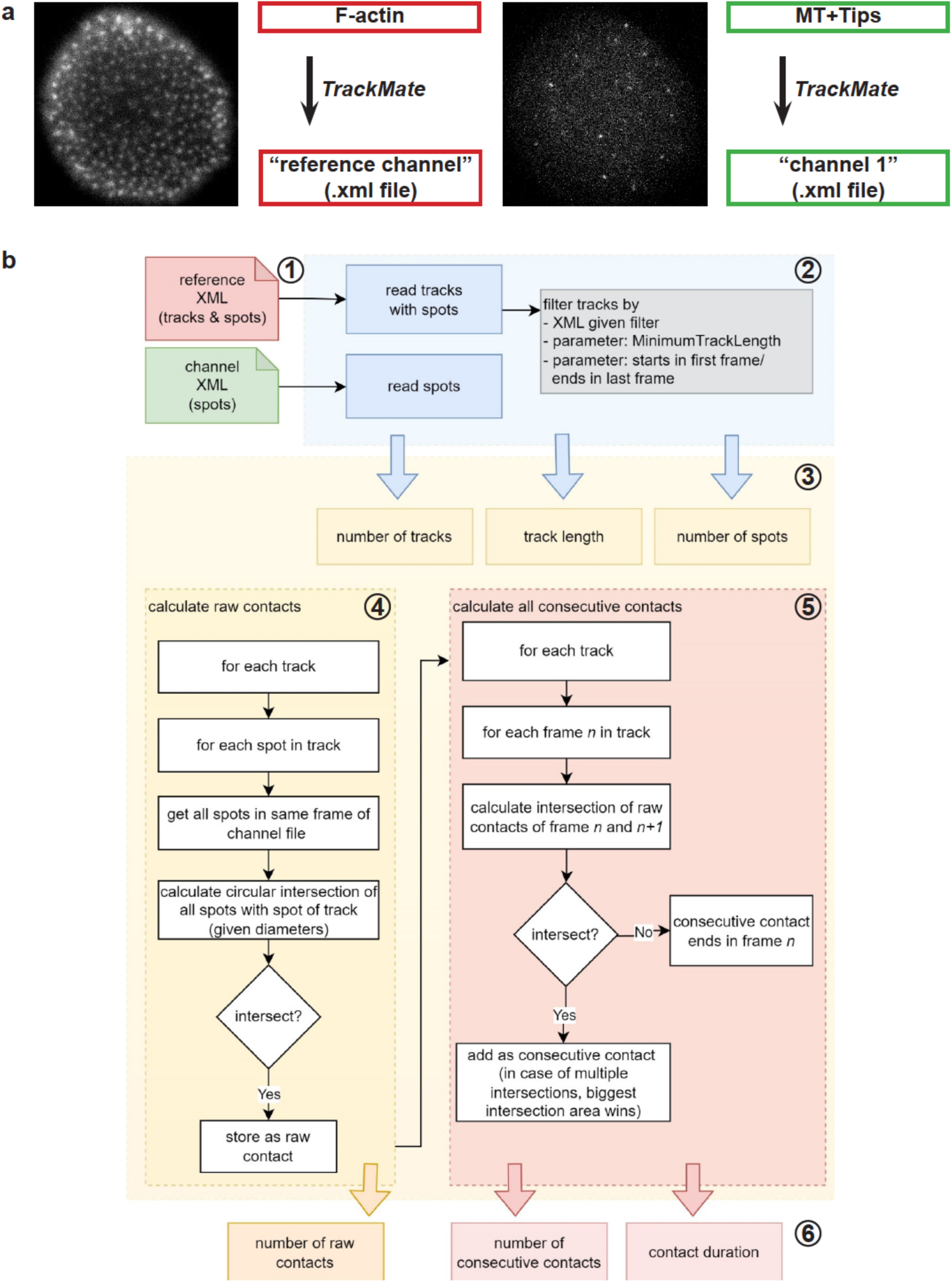
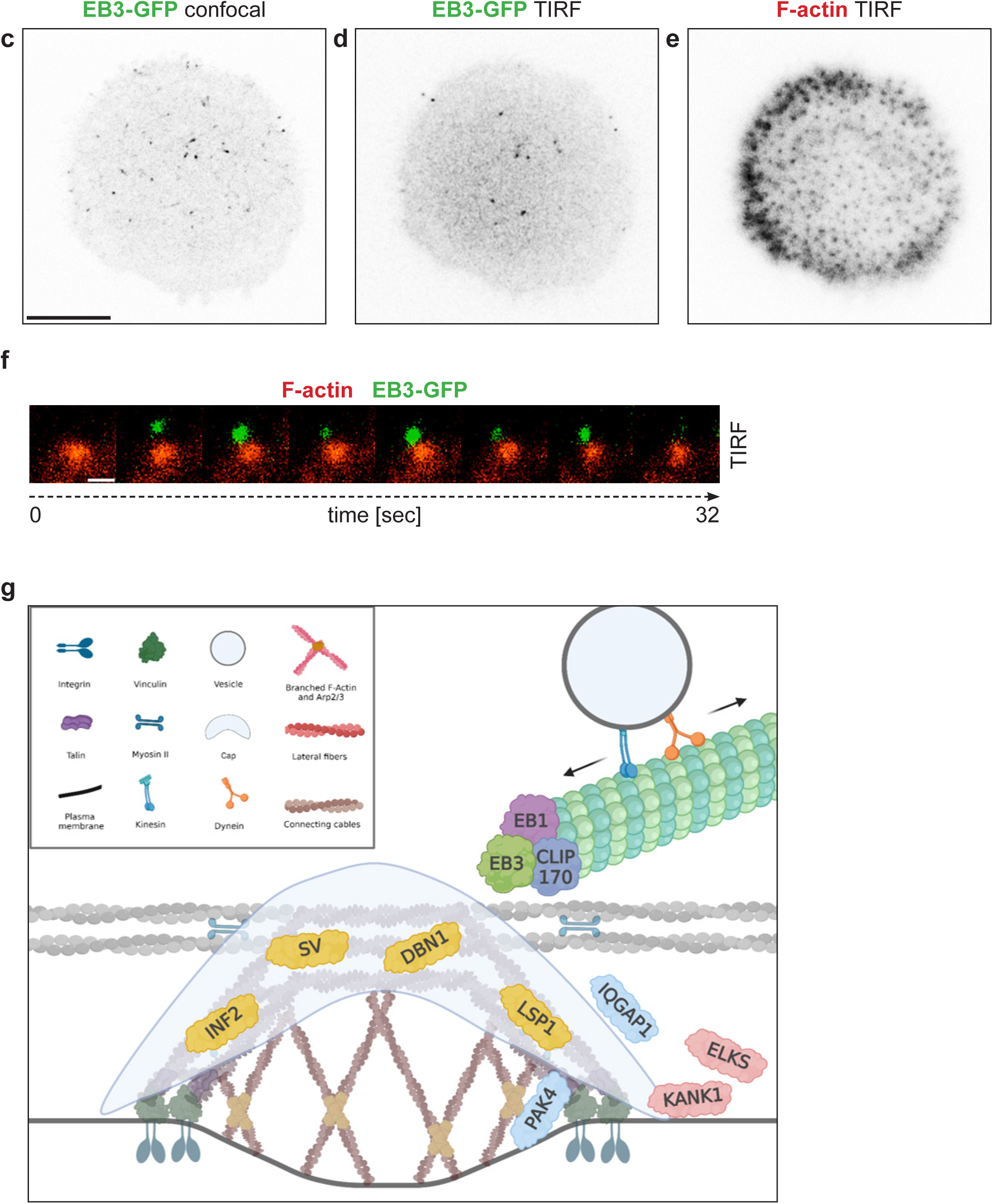
Analysis of contact between microtubule +tips and podosomes. (a) Acquisition of TIRF live cell videos showing podosomes, highlighted by overexpression of LifeAct-RFP (*reference channel*), and of TIRF live cell videos showing microtubule +tips, visualized by overexpression of EB3-GFP (*channel 1*). All movies are acquired with a frame rate of 0.5 frames per second. Podosome and +tip channel are subsequently opened in *ImageJ* and, using *TrackMate*, saved as .xml files with coordinates and tracks for podosomes and coordinates of +tips. (b) Work flow of the “*ContactAnalyzer*” macro. (1) XML files are generated from the acquired live cell videos (red symbol for tracks and spots, i.e. podosomes, and green symbol for spots, i.e. MT +tips); (2) Data files are read (blue box) and filters, parameters and coordinates are extracted and stay accessible (yellow boxes); (3) From this data pool, the number of tracks, their length and the number of spots are extracted; (4) All data are used in the first step to calculate the raw contacts; (5) In a subsequent step, the raw contacts are analyzed for being potentially consecutive; (6) In the end of the program run, result files are generated that contain information about the number of tracks and spots, the track length of podosomes, the number of raw and consecutive contacts, as well as the contact duration. (c-e) Micrographs of a primary human macrophage expressing EB3-GFP, highlighting microtubule +tips (c,d) and lifeact-RFP, highlighting podosome cores (d). EB3-GFP signals are detected in confocal mode (c) and TIRF mode (d), the latter highlighting only +tips that reach the podosome-containing ventral cell side. Respective signals shown in inverted black/white. Scale bar: 10 µm. (f) Contact event between a EB3-GFP positive +tip (green) and a lifeact-RFP-positive podosome (red). Detail region of TIRF video of cell shown in (d,e). Time is indicated in sec. Scale bar: 1 µm. (g) Model of podosome +tip contact with substructural localization of candidate proteins. Model shows podosome containing a core of branched F-actin (dark brown), surrounded by ring proteins that are connected through lateral unbranched actin filaments (grey) to the cap structure (yellow) on top of the core. A second set of unbranched actin cables (light brown), containing myosin IIA, connect individual podosomes. An incoming microtubule (green), featuring a +tip (blue) is depicted with a vesicle, transported by kinesin or dynein motors. Candidate proteins: cap components (yellow): INF2, supervillin (SV), LSP-1, drebrin (DBN1); actin-microtubule linkers (light blue): PAK4, IQGAP1; cortical docking complex components (pink): KANK, ELKS. Tested +tip proteins: CLIP-170 (dark blue), EB1 (purple), EB3 (green). Model created with Biorender.com.

To select a suitable marker for microtubule +tips, macrophages were transfected with GFP-fused constructs of CLIP-170, or of end binding protein 1 or 3 (EB1, EB3), and cotransfected with LifeAct-RFP, to visualize podosome cores (Ext. Data. Fig. 2a-c). Total interference reflection microscopy (TIRF) was chosen, as only +tips close to the ventral, podosome-containing cell side, were relevant for this analysis (Fig.1c-f), and 30 min videos (0.5 fps) of respective cells were analysed. Importantly, expression of all constructs was associated with comparable numbers of podosomes observed during duration of the videos (“cumulative podosome number”) (Ext. Data Fig. 2d). Differences were observed in all other parameters. The number of labelled +tips was lowest in cells overexpressing EB1-GFP, but comparable in cells expressing either of the other two constructs, with a trend towards higher numbers for cells expressing CLIP170-GFP (Ext. Data Fig. 2e). As podosome numbers were unchanged (Ext. Data Fig. 2d), a similar result was obtained for the number of raw contacts (Ext. Data Fig. 2f). The number of overall consecutive contacts, and also of consecutive contacts per podosome, was significantly lower for both EB1-GFP and EB3-GFP expressing cells, compared to those expressing CLIP-170-GFP (Ext. Data Fig. 2g,h). Regarding the duration of contacts, expression of EB1-GFP led to the shortest, and expression of EB3-GFP to the longest duration (Ext. Data Fig.2i). Based on the signal to noise ratio (Ext. Data 2a-c), and trying to avoid over-saturation of +tip labelling, EB3-GFP was chosen as a marker for subsequent experiments.

## Drebrin is a novel component of the podosome cap

In a screen for proteins regulating contact between podosomes and +tips, we included podosome cap proteins such as supervillin ^22^, INF2 ^26^, and LSP1 ^24^, and known regulators of +tip contact with focal adhesions, invadosomes, or cortical adhesions complexes, such as IQGAP1 ^50,51^, ELKS ^2,37^, and KANK1 ^2,38^ (Fig. 1g). We also included drebrin E2 ^52^ (hereafter called drebrin or DBN1), which we identified as a new cap component, both as an endogenous protein (Fig. 2a-c) and as a fluorescently labelled fusion construct (drebrin-GFP) (Fig. 2d-f). Poji macro ^53^-based analysis showed that drebrin exhibited a typical cap-like localization, with two maxima adjacent to the core-localized maximum of F-actin intensity in all analysed optical planes (Fig.2g-j) ^54^. This was confirmed by 3D reconstruction of single podosomes (Fig. 2k-n), while a comparison with the cap component α-actinin shows that drebrin has a broader distribution on both the xy and the xz axis and thus localizes to a more peripheral part of the cap (Fig. 2i,j).

**Figure 2.**
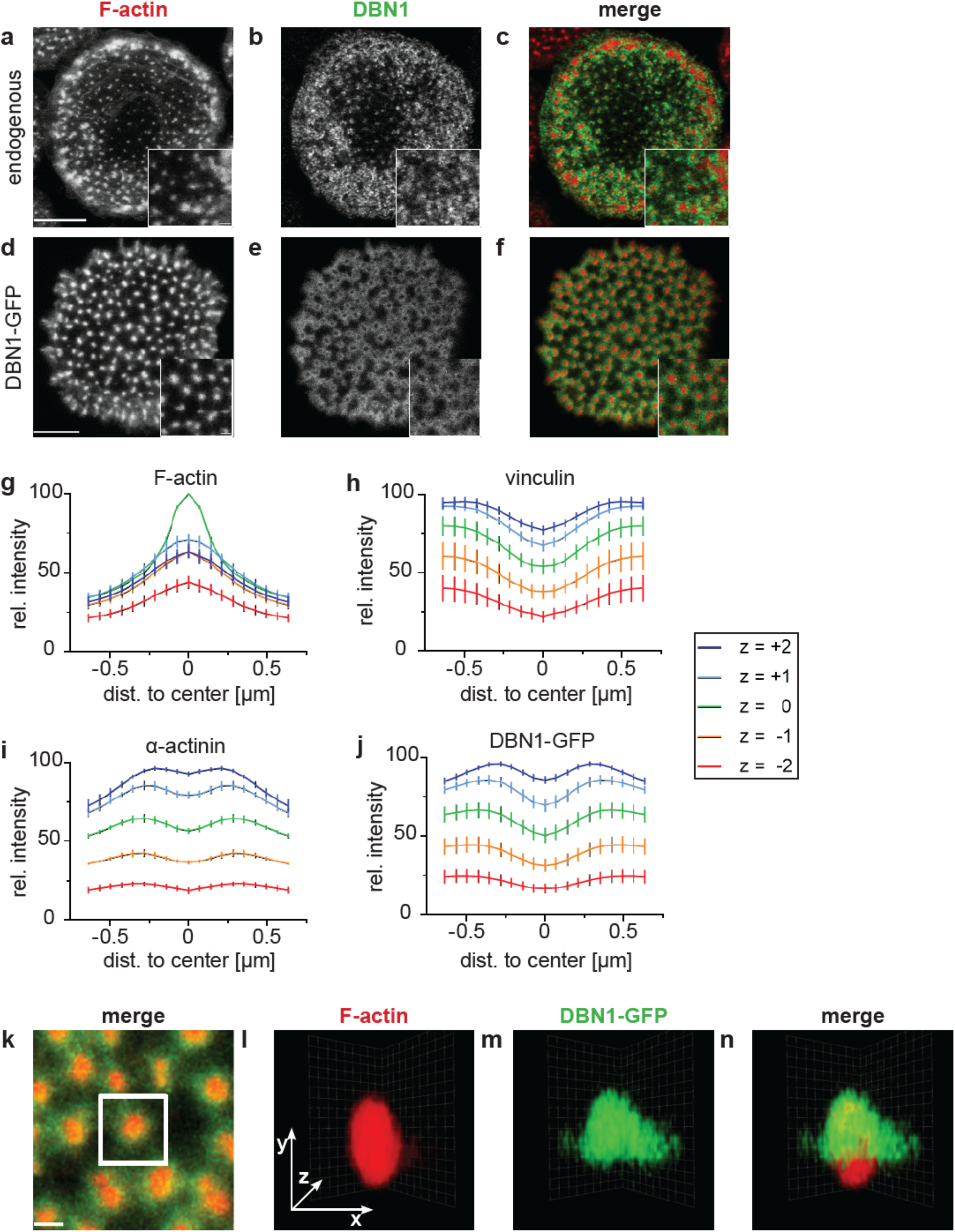
Drebrin is a component of the podosome cap. (a-d) Primary human macrophages stained for F-actin using Alexa 568-phalloidin (a,d), and for endogenous drebrin (b) or overexpressing drebrin-GFP (e), with merges (c,f). White boxes in (a,d) indicate regions of detail images in lower right. (g-j) Fluorescence intensity analysis at specific Z-planes of podosome core-localized F-actin (g), ring component vinculin (h), cap components α-actinin (i) and drebrin (j). Fluorescence intensity of proteins at podosomes was analysed using Poji macro in several optical planes with a distance of 250 nm each. Plane with maximal intensity for F-actin is marked as z=0, with colour code for optical planes on the right (n= 250 cells of 3 donors). (k-n) 3D rendering of a single podosome (n) reconstructed from the optical z-stack shown in the white box (k), with single channels showing F-actin (l) and DBN1-GFP (m) respectively. Scale bar: 10 µm for (a-f), 2 µm for (k).

Drebrin contains an N-terminal ADF homology region (ADFH; aa 1-135), followed by a coiled coil (CC) region regulating homodimerization (aa 176-256) ^55^, a helical domain (Hel; aa 257-355), a polyproline region (PP; aa 364-417) shown to bind to the endothelial adherens junction protein afadin ^52^, and a long unstructured C-terminal domain (Ext. Data Fig. 3a). Of note, the CC and Hel regions contain a “cryptic domain” that is able to independently bind to F-actin, but also to cooperatively bundle actin filaments ^48^. In order to identify the domains of drebrin that mediate its localization to podosomes, GFP-fused deletion constructs were overexpressed in primary macrophages and analysed by fluorescence microscopy, including a construct deleted for the ADFH region (ΔADFH-GFP), a construct deleted for the ADFH and polyproline regions (ΔADFH-ΔPP-GFP), as well as a construct containing only the CC and Hel regions (CC-Hel-GFP), a construct containing only the PP and C-terminal regions (PP-C-term-GFP), and a construct containing only the C-terminus (C-term-GFP). The ΔADFH-GFP and ΔADFH-ΔPP-GFP constructs showed a cap-like localization at podosomes (Ext. Data Fig. 3b,c), slightly more narrow to the F-actin core when compared to the full length protein (Fig. 2d-f), while the CC-Hel construct co-localized with the podosome core (Ext. Data Fig. 3d). Both the PP-C-term and the C-term-GFP construct showed a mostly dispersed localization (Ext. Data 3e,f).

We conclude that the CC, Hel and C-terminal regions together are required for localizing drebrin at the podosome cap substructure, whereas the ADFH and PP regions determine the degree of cap broadness, likely by regulating the amount of crosslinked or bundled actin filaments, as suggested by the current model of the podosome cap ^25^.

### Podosome cap proteins regulate different aspects of podosome/+tip contacts

SiRNA-based knockdowns using published sequences of candidate proteins (see Fig. 1g) resulted in remaining protein levels of 43%-0.2%, following normalization with GAPDH levels (Ext. Data Fig. 4a). Cells treated with specific siRNA or control siRNA and expressing EB3-GFP, to highlight +tips, and Lifeact-RFP, to detect podosomes, were visualized by live cell imaging in TIRF mode, and ten parameters for podosomes and +tips were extracted (see below).

The analysis showed that both i) cell area and ii) the number of podosomes per cell were reduced upon INF2 depletion (Ext. Data Fig. 4b,c), consistent with earlier results (Panzer et al., 2016). iii). The life time of podosomes was reduced in cells depleted for KANK1 (Fig. 3a), while iv) the cumulative number of podosomes over time were not significantly changed for all treatments (Fig. 3b). Furthermore, v) podosome density was significantly reduced in cells depleted for KANK1 and slightly increased for LSP1 depleted cells (Ext. Data Fig 4d). vi) The number of +tips per cell was slightly, but significantly, enhanced for IQGAP1-depleted cells, and decreased for drebrin-depleted cells (Ext. Data Fig. 4e). Strikingly, both vii) the number of raw contacts and of viii) consecutive contacts between podosomes and +tips, as well as ix) the number of contacts per podosome, was strongly decreased for drebrin-depleted cells (Fig. 3c-e), while x) the duration of contacts was increased for INF2-depleted cells and decreased for LSP1-depleted cells (Fig. 3f). Interestingly, a more detailed analysis of podosome life time showed that cells depleted for KANK1 or INF2 showed no longer lived (20 min – 30 min) podosomes. A similar trend was seen in cells depleted for supervillin or drebrin, where maximal podosome live times did not exceed 24 min or 28 min, respectively (Ext. Data Fig. 4f). Collectively, these results identify drebrin as a positive, and INF2 as a negative regulator of contact events between podosomes and +tips. In addition, INF2 emerges as a negative, and LSP1 as a positive regulator of contact duration.

**Figure 3.**
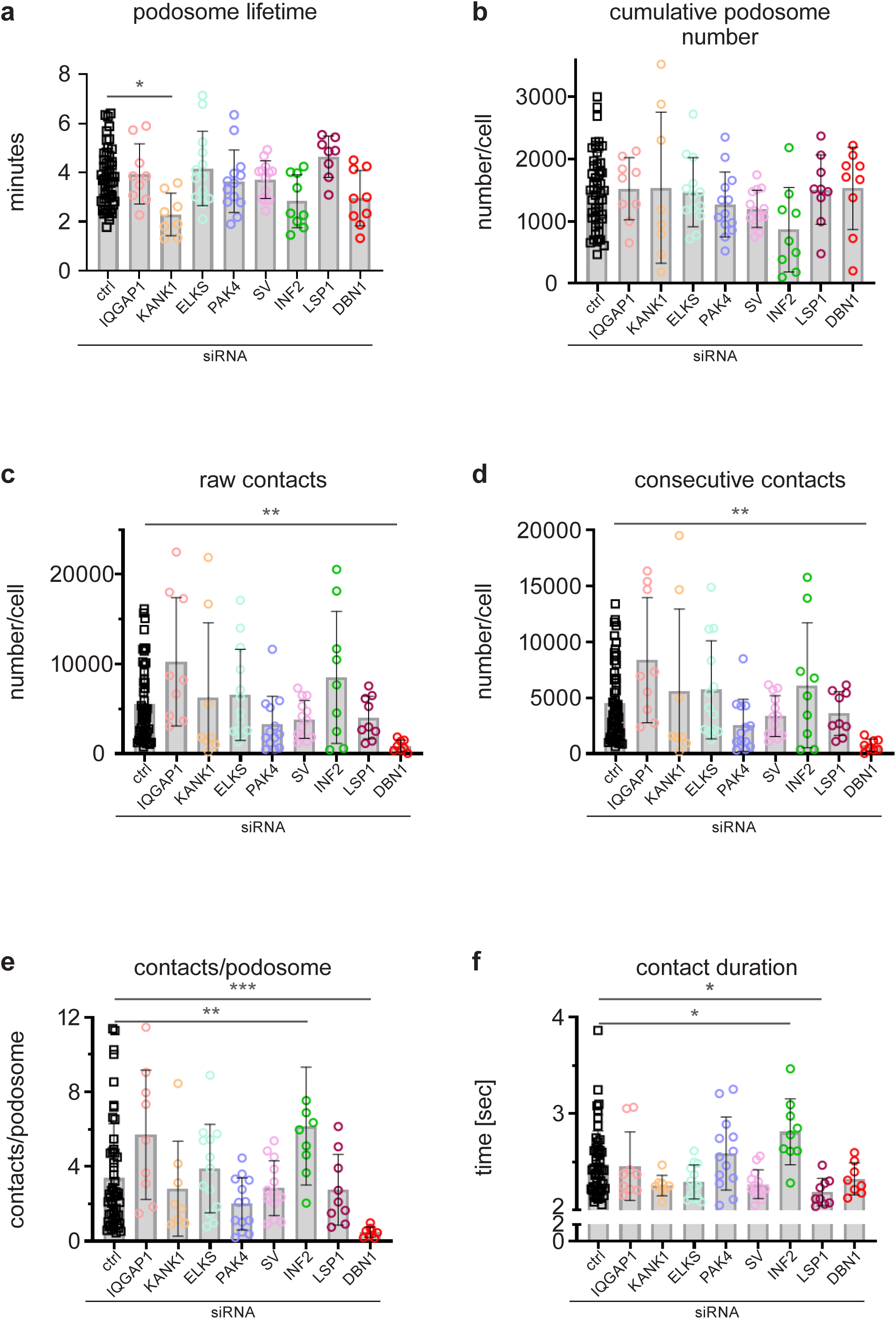
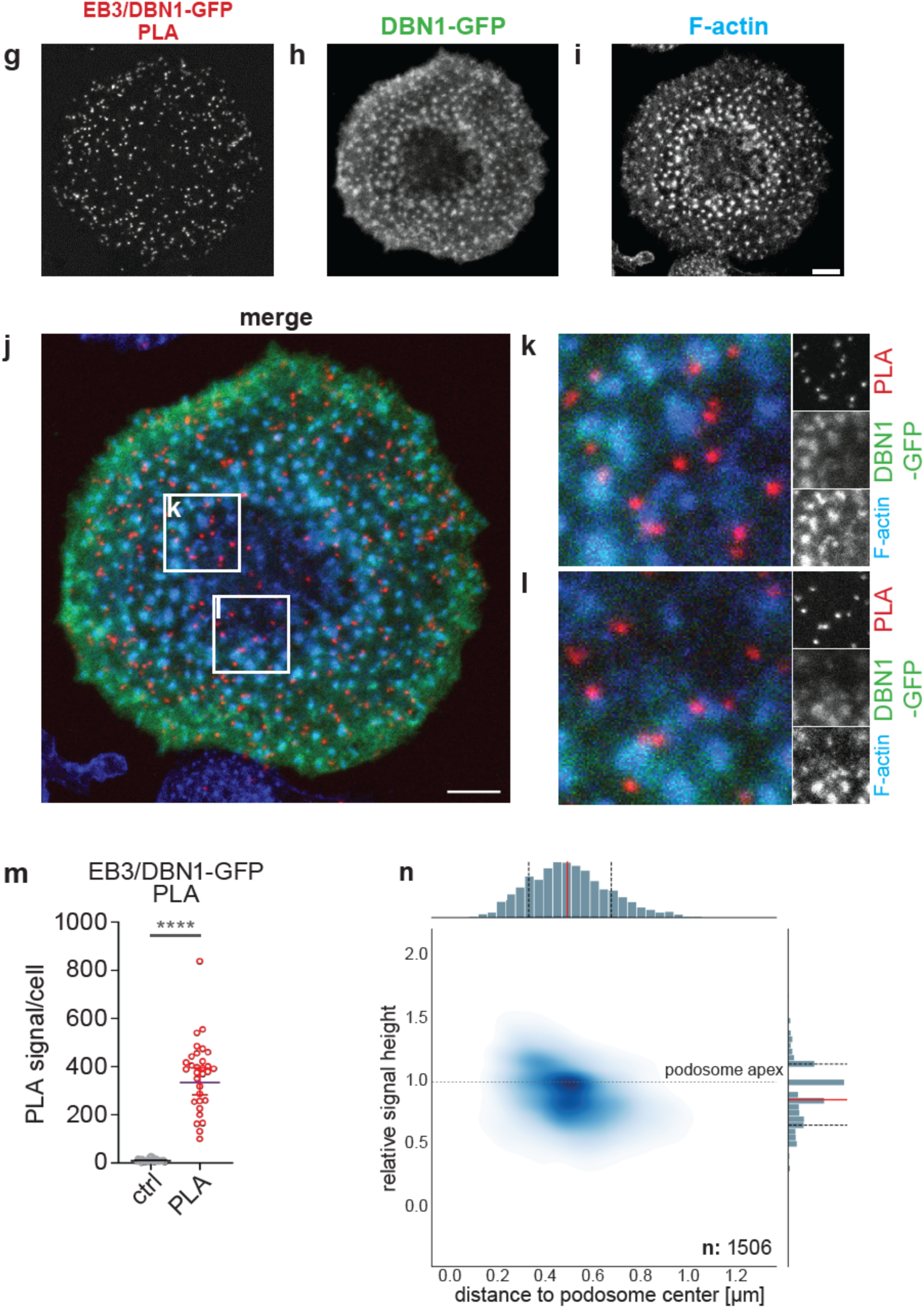
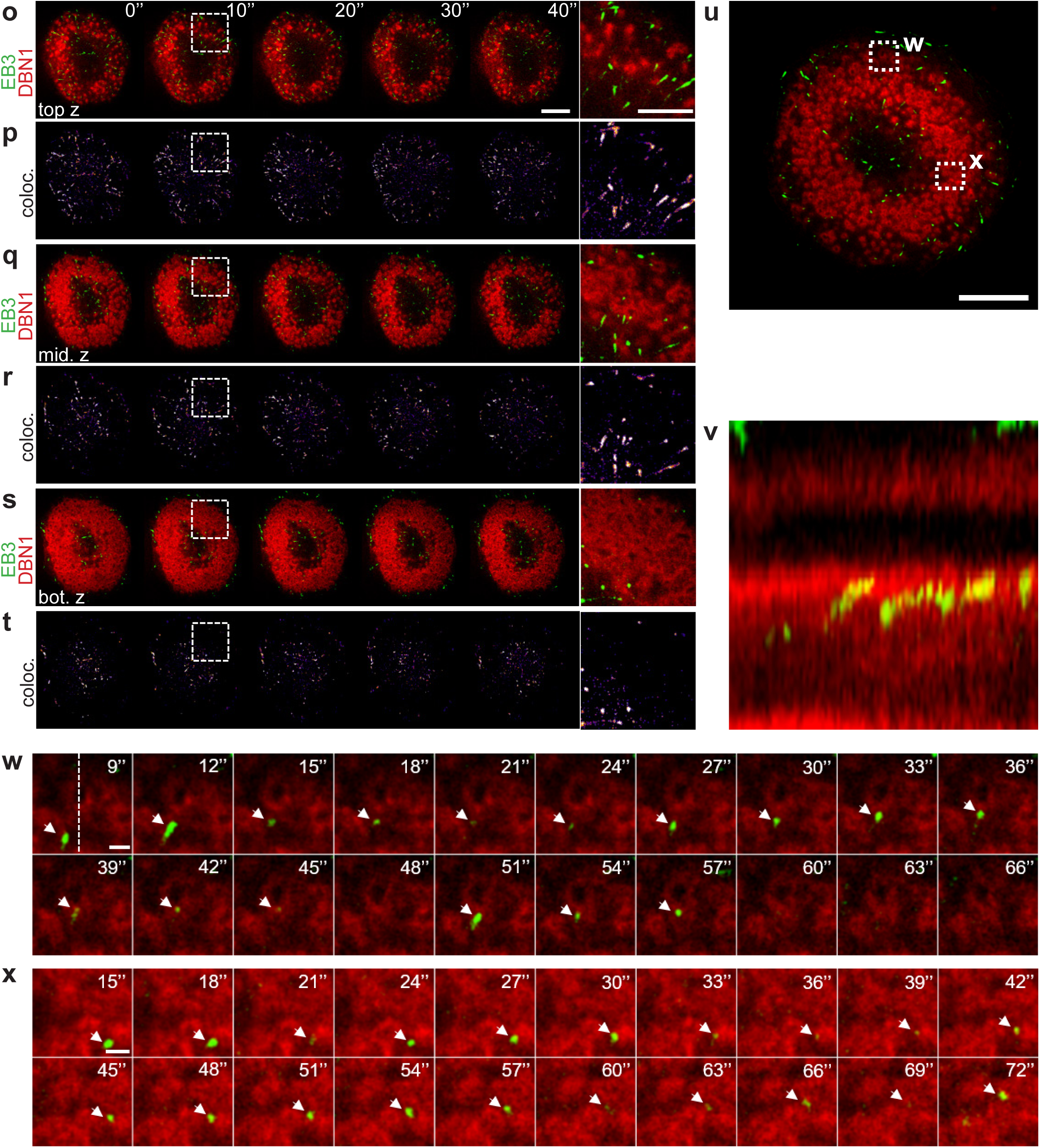
Podosome cap proteins differentially regulate contact with microtubule +tips; drebrin and EB3 share a close spatiotemporal relationship at podosomes. (a-f) Evaluation of podosome/+tip contacts during 30 min TIRF videos, using the ContactAnalyzer tool. (a) podosome lifetime, given in min, (b) number of all podosomes per cell observed during the 30 min videos (“cumulative podosome number”), (c) number of all contacts per cell observed during the 30 min videos (“raw contacts”), (d) number of consecutive contacts per cell, (e) number of contacts per podosome, (f) contact duration, given in sec. Statistics: non-parametric Kruskal-Wallis test, followed by Dunn’s multiple comparison post hoc test, \**P*<0.05, \*\**P*<0.01, \*\*\*\**P*<0.001. For all values, ≥ 8 videos of cells from ≥ 3 donors were analysed. For specific values, see Suppl. Table 2. (g-l) Confocal micrographs of macrophage processed for proximity ligation assay (PLA) using GFP- and EB3-specific antibodies, with respective PLA signals (g), with cell overexpressing drebrin-GFP (h), and stained with Alexa405-phalloidin for F-actin, to label podosome cores (i). White boxes in merge (j) indicate detail regions shown in (k,l), with respective single channels on the right. Note PLA signals adjacent to podosomes, but absent in podosome-free areas. Bars: 5 µm in overviews, 2 µm in detail images. (m) Statistical evaluation of PLA signals, compared to isotype controls. N= 3x10 cells, from 3 donors. ****P<0.001, according to Mann-Whitney test. (n) Spatial relationship between drebrin-EB3 PLA signals and podosomes. Kernel density estimation (KDE) plot with x-axis representing distance to podosome centroid in µm and y-axis showing relative signal height to the podosome apex. Frequency distributions of both measurements are shown along their respective axes. Gray horizontal line demarks the podosome apex. Distance to podosome centroid has a median of 0.49 µm with standard deviation of 0.17 µm. Relative height has a median of 0.86 with standard deviation of 0.24. n = 1,506 podosomes. Only podosome-associated signals were considered, which are signals in a 1 µm radius around a 3D podosome. N= 3x 10 cells from 3 donors. (o-x) EB3-GFP-positive MT +tips contact drebrin-RFP-positive podosome caps. Still images from 4D-SoRa time lapse videos of macrophage overexpressing Drebrin-RFP (red) together with EB3-GFP (green). (o-t) Galleries show top Z-slice (o) middle Z-slice (q), and bottom Z-slice (s), with colocalizing pixels shown in (p,r,t). Dashed white boxes indicate detail region shown on the right. Time is indicated in sec. Bars: 10 µm for overviews, 2 µm for ROIs. Note how most contacts between EB3 and podosomes occur in the middle-to-top part of podosomes. See Suppl. video 1. (u-x) Macrophage overexpressing drebrin-RFP (red) together with EB3-GFP (green); dashed white boxes indicate detail regions shown in (w,x). Bars: 10 µm for overview, 1 µm for ROIs, time is indicated in sec. (v) kymograph of pixels indicated by dashed line in (w). See also Suppl. videos 2-4.

**Figure 4.**
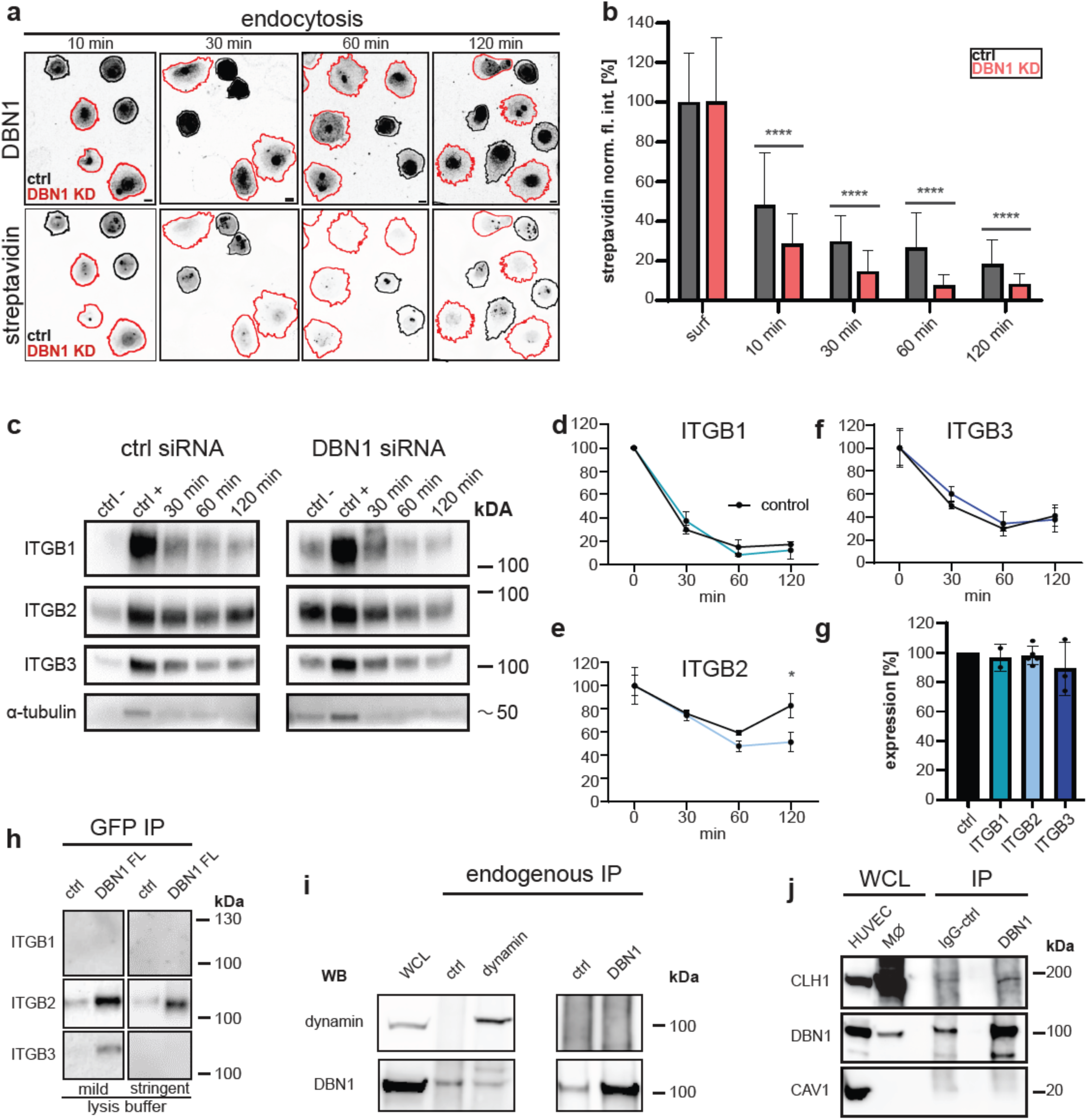
Drebrin depletion leads to reduced endocytosis, particularly of β2 integrin; drebrin does not interact with dynamin, but with clathrin heavy chain. (a) Time series of micrographs of primary human macrophages treated with control siRNA or drebrin siRNA, incubated with biotin to label surface proteins and with AlexaFluor488-streptavidin to visualize and quantify endocytosis of biotinylated proteins. Black outlines indicate cells with a high expression of drebrin, while red outlines mark cells with low expression. Note reduced streptavidin signal intensity (i.e. endocytosed surface proteins) in cells showing low drebrin expression and vice versa. (b) Quantification of streptavidin signal intensity, during endocytosis, in cells treated with control siRNA or drebrin siRNA. Signals are normalized to the mean intensity signal of the respective control. N= 49 cells from 4 independent donors, ****P<0.001, according to 2way ANOVA test. Bar: 10 µm. (c) Endocytosis of biotinylated surface proteins from cells treated with control siRNA or drebrin siRNA with representative Western blots showing β1, β2, β3 integrins, and α-tubulin as loading control. Time since start of the experiment is indicated in min. (d-f) Evaluation of signal intensities of blots from (c) Signals are normalized to mean intensity of drebrin signal of respective control. N= 2 independent donors, *P<0.05, according to multiple unpaired T-tests. (g) Analysis of total cellular expression levels of target proteins in primary human macrophages treated with drebrin siRNA for 72 hrs: N= at least 2 samples from independent donors; ns, according to One-way ANOVA test. (h) Western blot of anti-GFP immunoprecipitations from lysates of macrophages expressing drebrin-GFP or GFP empty vector as control, lysed with mild or stringent lysis buffers, and developed with anti-β1, β2, or β3 integrin antibodies. Note prominent coprecipitation of β2 integrin in both conditions and faint coprecipitation of β3 integrin only in mild condition. (i) Western blot analysis of cross-immunoprecipitations of endogenous dynamin or drebrin from macrophage lysates, with whole cell lysate (WCL) as control. Note absence of drebrin in anti-dynamin IP and vice versa. (j) Western blot analysis of whole cell lysates from HUVEC or primary human macrophages (left), and immunopreciptations from human macrophage lysates using control IgG or anti-drebrin antibody, probed for clathrin heavy chain, drebrin, and caveolin-1. Note detection of caveolin-1 in lysates from HUVEC, but not from macrophages. Also note co-precipitation of clathrin heavy chain with drebrin. Molecular weight in kDa is indicated on the right for (c,h-j).

Identification of cap-localized drebrin as a regulator of contact with EB3-positive +tips is in agreement with previous findings showing that drebrin, together with EB3, regulates contact of +tips with the cell cortex of rat embryonal cortical neurons ^56^. However, while the direct binding between drebrin and EB3 is well established ^49,57,58^, it is unclear at which subcellular sites this interaction takes place. We therefore performed proximity labelling assays (PLA) using primary macrophages overexpressing drebrin-GFP, together with anti-EB3 and anti-GFP antibodies, and staining with Alexa568-phalloidin to visualize podosomes (Note: overexpression of drebrin-GFP was chosen as primary antibodies against drebrin and EB3 were not compatible for PLA). Respective cells showed numerous PLA signals (Fig. 3g-m), which were largely absent in isotype controls (Fig. 3m). Of note, PLA signals were adjacent to podosomes (Fig. 3k), while podosome-free areas did mostly not show PLA signals (Fig. 3l). Drebrin and EB3 thus share a close spatial relationship (<40 nm) at podosomes. Use of a newly developed algorithm (“BioPixel”, see Materials and Methods) further showed that PLA signals have a median distance of 0.49 µm from the centroid of podosomes in the xy plane, and a median distance of 0.86 µm from the podosome bottom in the xy plane (Fig. 3n). These measurements indicate that respective PLA signals are directly adjacent to podosomes, and also near to their apex, as podosomes have a typical diameter and height of ∼1µm ^59^. This was further confirmed by colocalization analysis of live cell videos showing that drebrin-RFP and EB3-GFP were often in a close spatial relationship at the top, and also at the side of podosomes (Fig. 3o-t). Further analysis showed that EB3-GFP positive +tips dynamically and repeatedly contacted drebrin-RFP positive podosomes (Fig. 3u-x).

### Drebrin regulates endocytosis of β2 integrin

The connection between drebrin at the podosome cap and EB3 at MT +tips indicated that drebrin could be involved in the regulation of transport events at podosomes. In combination with an earlier study implicating drebrin in endocytosis of β1 integrin in adenocarcinoma cells ^60^, this prompted us to investigate a potential role of drebrin in endocytosis at podosomes. A generalized endocytosis assay using cell-impermeable, cleavable biotin to label cell surface proteins, and streptavidin as a marker, showed that low levels of drebrin in respective knockdown macrophages correlated with a continuous decline of intensities of endocytosed streptavidin (i.e. biotinylated surface proteins) (Fig. 4 a,b). Integrins such as β1- ^61^, β2- ^61^, and β3 integrin ^42^ are prominent podosome-associated transmembrane proteins. To investigate a potential influence of drebrin on integrin endocytosis, we measured cell surface levels of β1, β2, and β3 integrin in a standardized surface biotinylation assay. Lysates of siRNA treated cells were collected at timepoints 0, 30, 60 and 120 minutes after the initial labelling with cell-impermeable, cleavable biotin. Western blot analysis of the respective lysates showed that the levels of endocytosed β1 and β3 integrin, were not significantly changed upon drebrin knockdown. In contrast, the amount of endocytosed β2 integrin was significantly altered upon the depletion (Fig. 4c-e), while total cellular expression levels of all investigated proteins were unchanged (Fig. 4g). Collectively, these data pointed to a role of the drebrin in endocytosis of β2 integrin at podosomes. We next investigated a potential interaction between drebrin and β1-3 integrins using anti-GFP immunoprecipitation in lysates of macrophages overexpressing drebrin-GFP. Using a mild lysis buffer, β2 integrin and, faintly, also β3 integrin could be coprecipitated with drebrin-GFP. Using a more stringent buffer, only β2 integrin was coprecipitated. In both cases, β1 integrin was not detected in respective precipitates (Fig. 4h). We next investigated whether the apparent specificity of drebrin-dependent endocytosis for β2 integrin could be due to its localization at podosomes. Indeed, both confocal and TIRF microscopy showed a prominent localization of β2 integrin at the podosome ring structure, while β3 integrin was mostly detected at the cell periphery, with moderate localization at podosome ring, and β1 integrin was only poorly colocalizing with podosomes (Ext. Data Fig. 5).

As drebrin was previously reported to interact with the endocytosis-associated GTPase dynamin ^60^, we performed cross-immunoprecipitations for both proteins in their endogenous forms. However, no copreciptation above background control levels was discernible for either protein (Fig. 4i). We next investigated potential further connections of drebrin to the endocytic machinery. Of note, β2 integrin endocytosis has been mostly studied in the context of clathrin-dependent pathways^62^ ^63^. Moreover, drebrin does not feature a caveolin-interaction motif, and in silico analysis of a potential interaction between drebrin and caveolin-1 using AlphaFold3 yielded only a low score (ipTM: 0.12). Furthermore, GeneCards annotation (www.genecards.org) showed that caveolin-1 and -2 are not expressed in human monocytes, which was also confirmed by respective Western blots of lysates from human macrophages (Fig. 4j). Accordingly, clathrin heavy chain (CLH1), but not caveolin-1, could be co-immunoprecipitated with drebrin from macrophage lysates (Fig. 4j)

### Drebrin binds clathrin heavy chain through a canonical and a novel clathrin box

We next focused on a potential connection between drebrin and clathrin. Immunofluorescence analysis revealed numerous clathrin heavy chain (CLH1)-positive signals on the ventral cell macrophage surface that were often in close proximity to podosomes (Fig. 5a-h), while colocalization analysis showed numerous overlaps between drebrin and CLH1 at podosomes (Fig. 5i,j). Proximity ligation assays using antibodies specific for drebrin and CLH1 showed numerous PLA signals (Fig. 5k-p) that were mostly adjacent to podosomes and absent from podosome-free areas (Fig. 5n,o). BioPixel analysis revealed that respective PLA signals had a median distance of 0.48 µm from the centroid of podosomes in the xy plane, and a median distance of 1.0 µm from the podosome bottom in the xy plane (Fig. 5q), indicating that respective PLA signals were directly adjacent to podosomes and near to their apex (compare to Fig. 3n). This was confirmed by staining of endogenous CLH1 and EEA1, which showed numerous colocalizing signals at the side or the top of the podosome cap, visualized by overexpression of drebrin-GFP (Fig. 6r-w2), Furthermore, live cell imaging of macrophages overexpressing clathrin light chain (CLL1)-GFP and drebrin-RFP revealed numerous dynamic contacts between clathrin vesicles and podosomes (Fig. 5x), with clathrin vesicles sometimes emerging directly adjacent to podosomes (Fig. 5x1). Colocalization analysis further showed that clathrin vesicles contacted podosomes especially during podosome growth and dissolution (Fig. 5x2).

**Figure 5:**
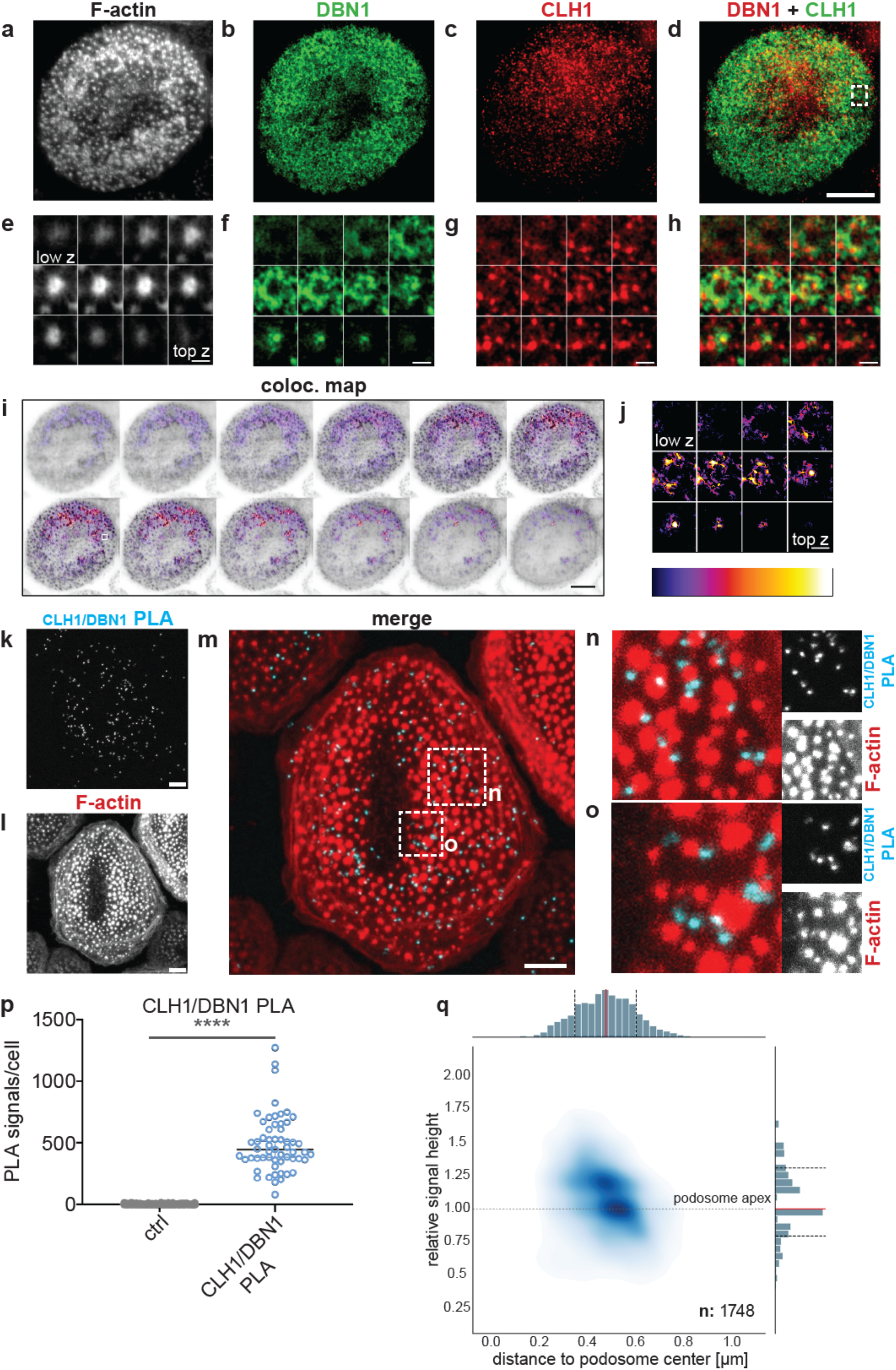
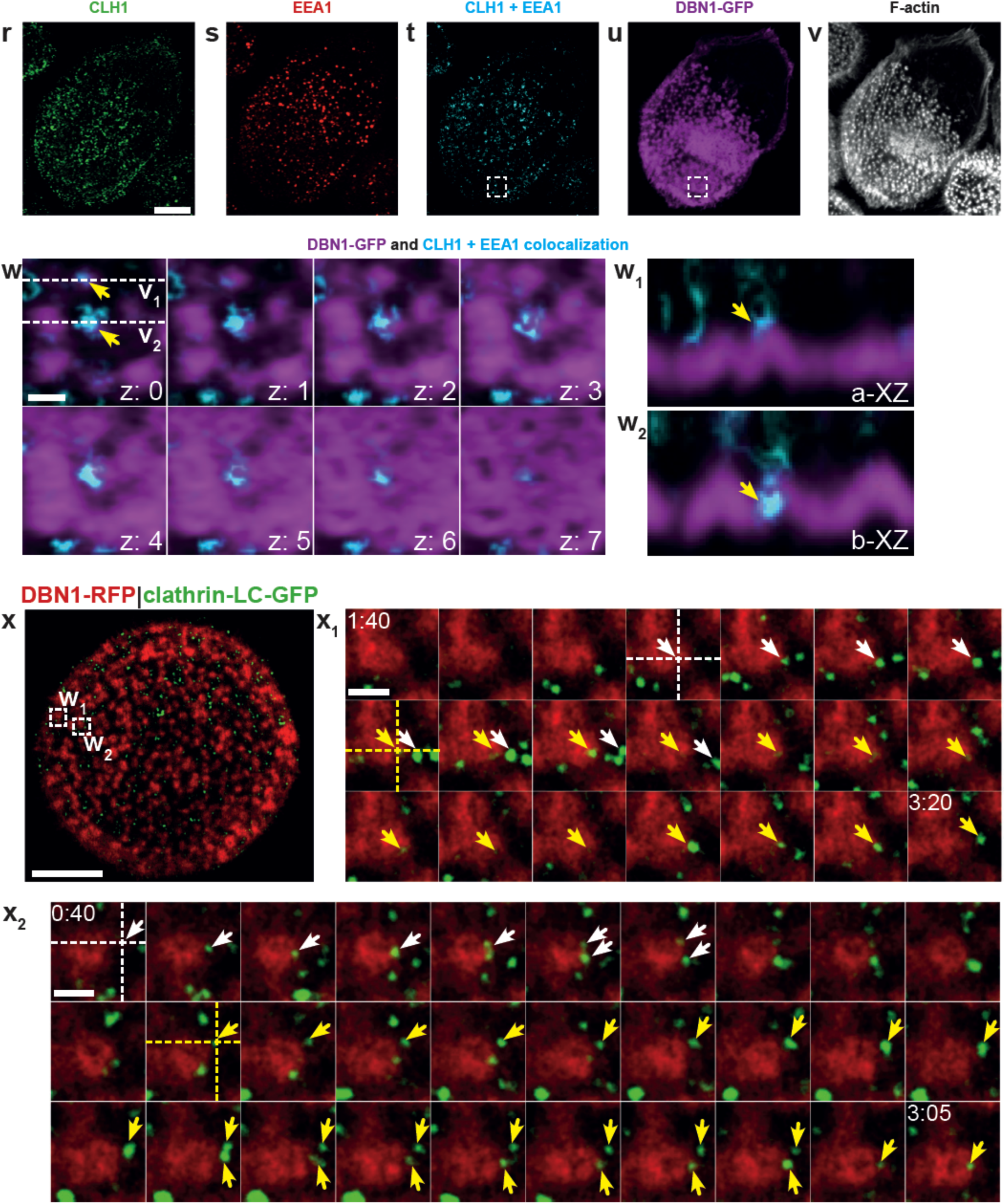
Drebrin and clathrin are closely associated at podosomes. (a-j) Confocal micrographs of primary macrophage stained for F-actin (a, white), drebrin (b, green), clathrin heavy chain (c, red), with merge (d). White dashed box (d) indicate area of detail regions show at different confocal planes in (e-h). Note partial colocalization of drebrin and clathrin heavy chain especially at outer layers of the podosome cap. Respective colocalization maps are shown as heatmap-colored images in (i,j). (k-p) Drebrin and clathrin are closely associated at podosomes. Confocal micrographs of macrophages, with single channel images showing (k) Proximity Ligation Assay (PLA) signal (blue in merge), using anti-drebrin and anti-clathrin heavy chain antibodies, (l) F-actin (red in merge), with merge (m). White boxes in (m) indicate regions of interest shown in (n,o). Note PLA signals in close proximity to podosomes, while being absent in podosome-poor regions. All images are maximum intensity projections from z-stacks. Bars: 5 µm for overviews, 2 µm for detail images. (p) Statistical evaluation of PLA signals, compared to isotype controls. N= 3x10 cells, from 3 donors. ****P<0.001, according to Mann-Whitney test. (q) Kernel density estimation (KDE) plot with x-axis representing distance to podosome centroid in µm and y-axis showing relative signal height to the podosome apex. Frequency distributions of both measurements are shown along their respective axes. Gray horizontal line demarks the podosome apex. Distance to podosome centroid has a median of 0.48 µm with standard deviation of 0.13 µm. Relative height has a median of 1.00 with standard deviation of 0.26. n = 1,748 podosomes. Only podosome-associated signals were considered, which are signals in a 1 µm radius around a 3D podosome. N=3x10 cells from 3 donors. (r-w) Clathrin heavy chain and EEA1 colocalize at the podosome cap. Maximum projections of confocal micrographs of primary human macrophages, stained with respective antibodies for endogenous CLH1 (r, green), EEA1 (s, red), with colocalizing pixels (t, cyan), and overexpressing drebrin-GFP (u, magenta), and stained with Alexa405-phalloidin for F-actin (v, white). Dashed white boxes in (t) and (u) indicate detail region shown in (w), with drebrin-GFP signal together with CLH1 and EEA1 colocalizing pixels. Different confocal planes are shown, starting from the top to the ventral side of podosomes. Dashed lines in (w) indicate xy views shown in (w1, w2). Yellow arrowheads indicate close association of drebrin-GFP and CLH1/EEA1 colocalizations at podosomes. See also Ext. Data Fig.8b-c showing single Z-planes. (x) Macrophage expressing drebrin-RFP and clathrin light chain (CLC)-GFP. Still image from live cell video. White boxes indicate detail regions shown as galleries in (x1,x2). White and yellow arrowheads indicate individual CLC-positive vesicles emerging near podosomes, with fission events in (x2). White and yellow dashed cross-sections indicate specific regions, proximal to podosomes, where CLC-GFP vesicles emerge consecutively. Time since start of experiment is indicated in min:sec. Bars: 10 µm in overviews, 1 µm in detail images. See Suppl. Videos 5-7.

**Figure 6:**
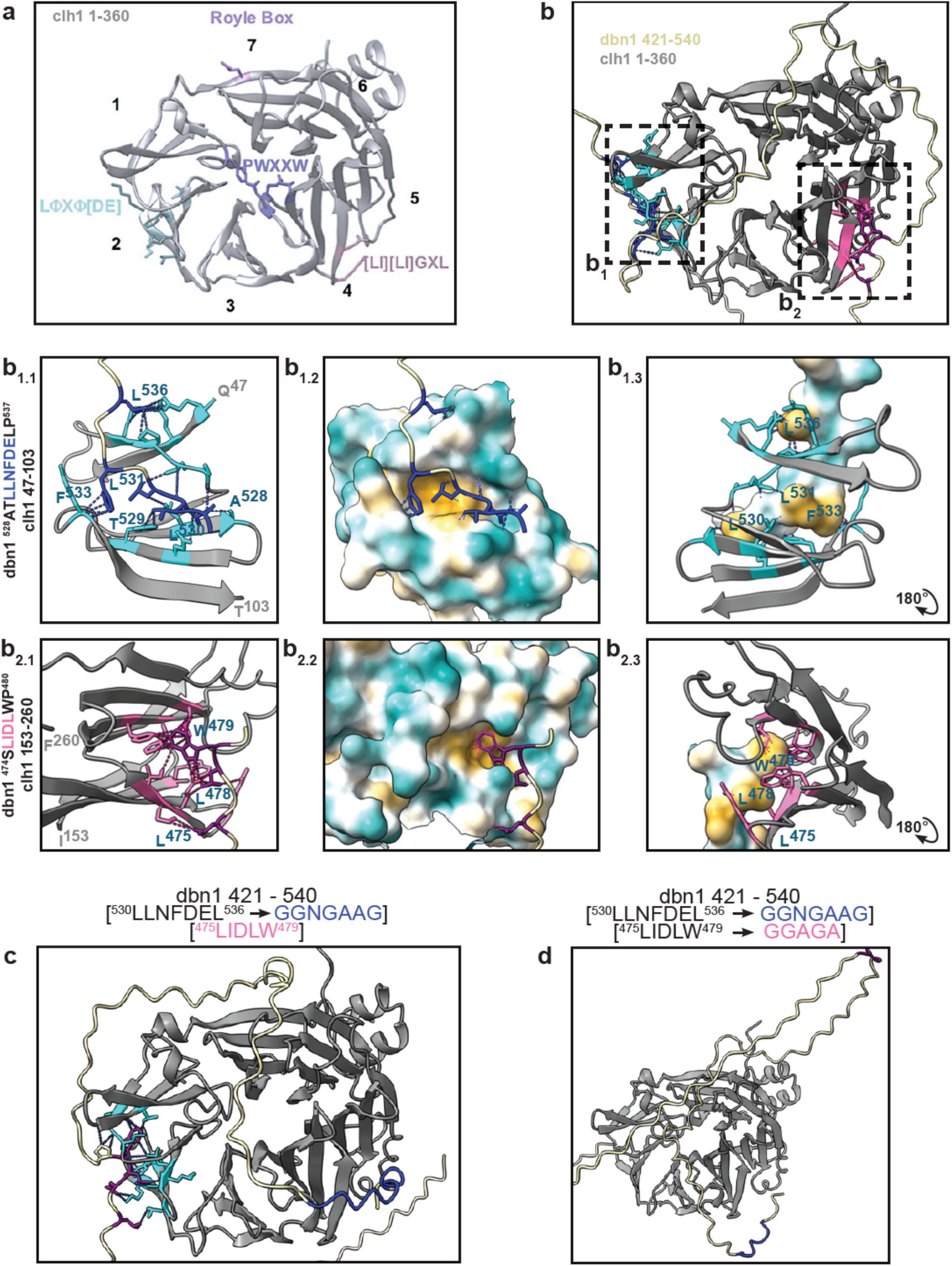
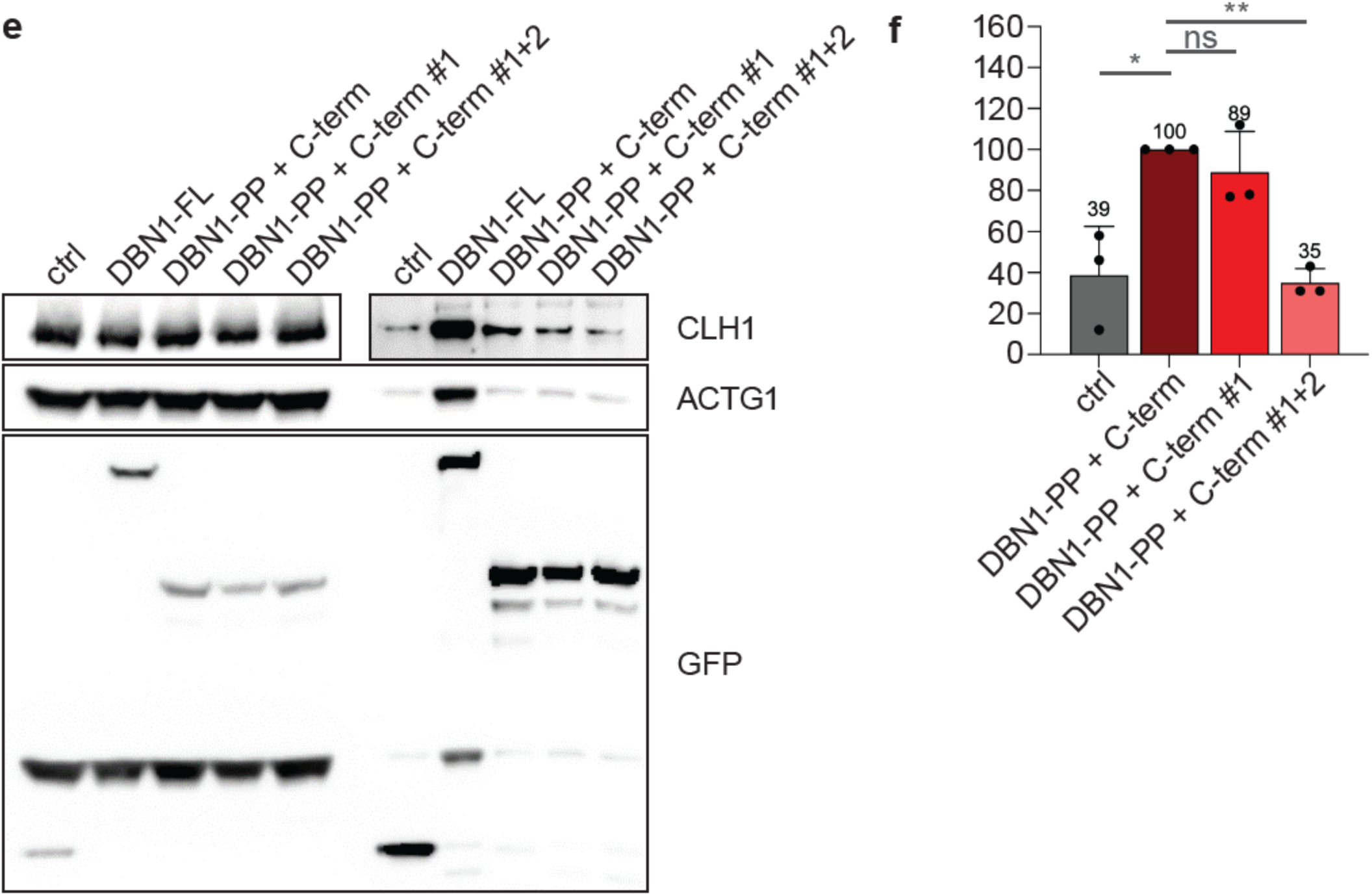
Analysis of CLH1 binding sites of the drebrin C-terminus. (a) Ribbon diagram of the CLH1 N-terminus, with numbered β-propeller domains, and with coloured regions showing the binding sites of interaction partners together with their respective motifs (boxes) involved in the binding, which include the canonical β-arrestin box (LΦx Φ[D/E]), the amphiphysin W-box ((P)WxxW), the arrestin box, ((L/I)(L/I)GxL), and the Royle box. Modified from ^95^. (b) AF3 model of the CLH1 N-terminus (aa1-360, gray) in contact with at least two motifs present in the C-terminus of drebrin (aa 421-540), with interacting amino acid residues of CLH1 (cyan and magenta) and drebrin (dark blue and purple). ipTM score of 0.72 and pTM score of 0.76. Dashed lines represents pseudobonds of interacting atoms with Van der Waals overlap ≥ - 0.1 [. Dashed boxes indicate detail regions shown enlarged in (b1.1, b2.1), with further surface rendering of CHL1 (yellow: hydrophobic; cyan: hydrophilic) (b1.2, b2.2), or of the respective drebrin motifs, rotated by 180° (b1.3, b2.3). (see also Ext. Data Fig. 6a,b for polar interactions). (c,d) AF3 models of mutations in the drebrin C-terminus that abrogate CLH1 binding. (c) Extended ^530^LLNFDEL^536^ motif mutated to GGNGAAG (blue); note relocalization of the ^475^LIDLW^479^ motif (magenta) to the site of the canonical clathrin box (cyan). (d) Additional mutation of the ^475^LIDLW^479^ motif to GGAGA; note complete loss of CLH1 binding by both motifs. (e,f) Anti-GFP immunoprecipitation from lysates of macrophages expressing indicated C-terminal drebrin constructs (wild type, mutated in box #1, or mutated in boxes #1+2; see text), with GFP as control; molecular weight in kDa indicated on the left. (e) Western blots showing co-precipitated CLH1 or ACTG1 (cytoplasmic γ-actin), and GFP control. (f) Quantification of co-precipitated CLH1 levels (a.u.), normalized to ACTG1, and with CLH1 levels in GFP controls set as baseline (n=3), *P<0.05, **P<0.01, according to Student’s t-test.

CLH1 contains four sites, also called boxes, in its N-terminal β-propeller domain that enable binding of interaction partners in hydrophobic pockets: i) a canonical box, represented by the sequence LΦx Φ[D/E] (where x refers to any amino acid and Φ to a bulky hydrophobic residue) ^64^, ii) a W-box that contains the (P)WxxW motif of amphiphysin ^65^, iii) the β arrestin box, which binds the (L/I)(L/I)GxL motif of arrestin 2 ^66^, iv) and the so called Royle box ^67^ (Fig. 6a). Computational analysis revealed that drebrin harbours a canonical clathrin box, represented by the sequence ^530^LLNFD^534^. Additionally, a further linear motif, ^475^LIDLWP^480^, shows partial similarity with the arrestin box, and, when inverted, also with the W-box. We then used AlphaFold3 to elucidate the putative binding sites of these motifs, in the context of the drebrin C-terminus (aa 421-540), to the CLH1 N-terminus (aa 1-360). Respective AF3 models place the ^530^LLNFD^534^ motif at the site of the canonical clathrin box, while the ^475^LIDLWP^480^ sequence is placed at the site of the arrestin box (iPTM=0.72) (Fig. 6b), with both sequences showing numerous amino acid residues potentially interacting with the CLH1 surface (Fig. 6b,1-1 to 2-3; Suppl. videos 5,6), based on complementary hydrophobicity (Fig. 6b) and electrostatic (Ext. Data Fig.6a-c) environments. We next used AF3 to identify mutations in these motifs that could lead to loss of CLH1 binding. Accordingly, mutation of the extended ^530^LLNFDEL^536^ motif to GGNGAAG led to loss of this sequence to bind the site of the canonical clathrin box. Instead, the ^475^LIDLWP^480^ motif was now modelled to bind this box, which could indicate a certain level of promiscuity of the drebrin boxes in binding different sites on the CLH1 N-terminus (Fig. 6c). Additional mutation of the ^475^LIDLW^479^ motif GGAGA led to complete loss of CLH1 binding by the drebrin C-terminus (Fig. 6d).

To test these predictions, we generated GFP-fused constructs of the C-terminal half of drebrin (GFP-PP-C-term; Ext. Data Fig.3) with mutations in the two potential CLH1 binding sites: one in which the extended ^530^LLNFDEL^536^ motif was mutated to GGNGAAG (GFP-drebrin-PP-C-term #1), and one in which also the ^475^LIDLW^479^ motif was mutated to GGAGA (GFP-drebrin-PP-C-term #1+2) (Note: Use of full length drebrin, including the N-terminal actin binding domains, showed substantial co-precipitation of actin, thereby masking potential interactions between drebrin and CLH1). Immunoprecipitations with wild type GFP-PP-C-term drebrin and GFP as controls, corrected for GFP-associated background levels, showed a ∼11% reduction in coprecipitation of CLH1 by the GFP-drebrin-PP-C-term #1 construct, and a significant reduction to, or slightly below, background levels by the GFP-drebrin-PP-C-term #1+2 mut construct (Fig.6e,f), thus validating the AF3 predictions, and showing that both sites on the drebrin C-terminus are necessary and sufficient for binding of CLH1.

Collectively, these data indicate that drebrin at the podosome cap represents a multivalent, actin-based hub that regulates endocytic trafficking of β2 integrin. Drebrin is able to associate with β2 integrin, and to bind CLH1 through the newly described clathrin binding boxes, thus linking the endocytic target with the endocytosis machinery. Moreover, binding of drebrin to EB3 allows access of β2 integrin-carrying clathrin vesicles to intracellular trafficking pathways through the microtubule system (Fig. 7a; see Discussion).

**Figure 7:**
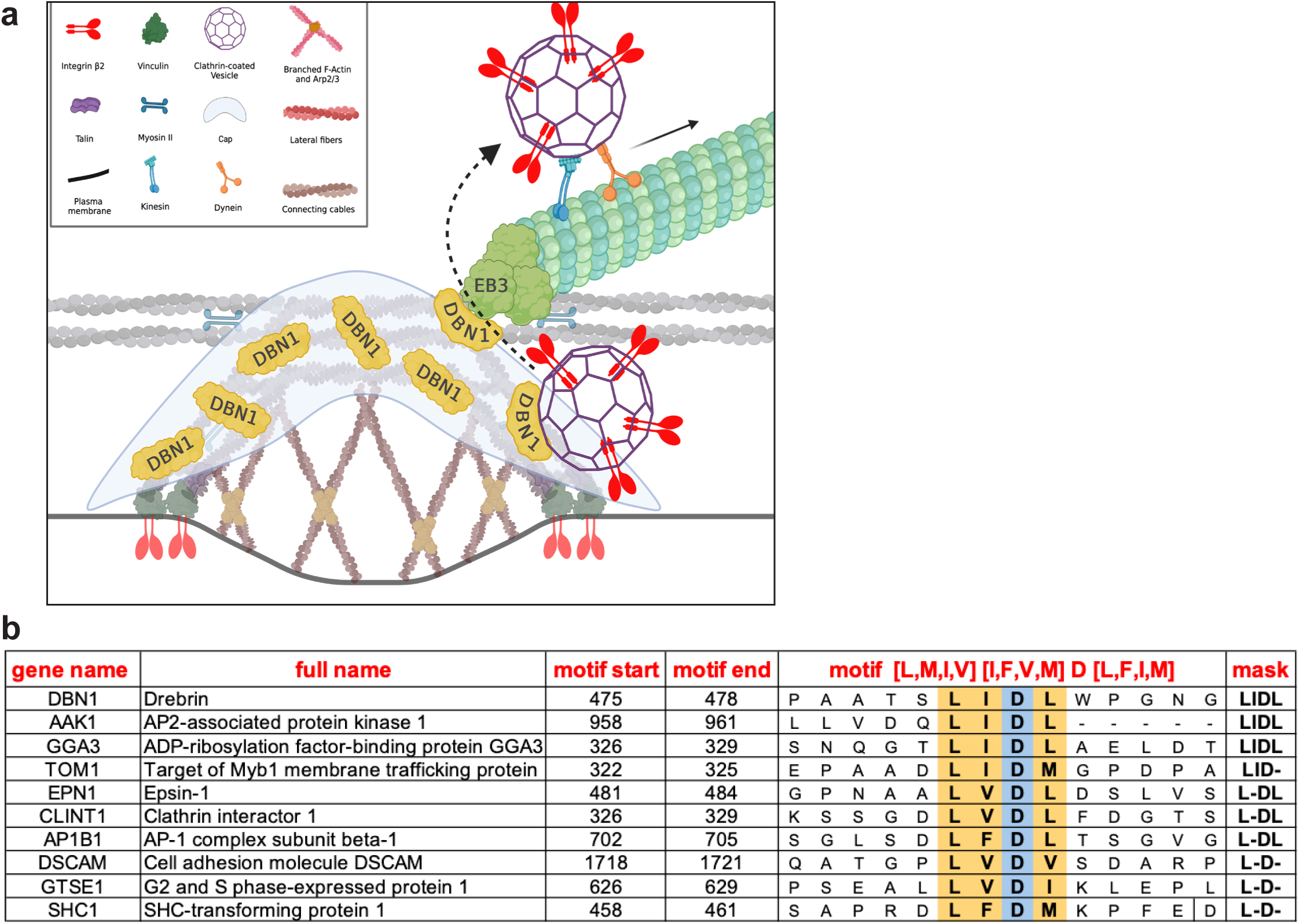
Model of drebrin as an endocytic hub for β2 integrin; occurrence of the LIDL motif in mammalian proteins. (a) Model: Microtubule +tips are connected to podosomes by binding of +tip localized EB3 to drebrin at the podosome cap. The drebrin C-terminus contains two CLH1 binding motifs, drebrin thus functions as a hub that links the actin cytoskeleton with clathrin-based endocytosis and microtubule-based transport (see also Fig. 1G). Model created with Biorender.com. (b) Bioinformatic analysis shows that the [L,M,I,V] [I,F,V,M] D [L,F,I,M] motif is also present in a variety of other mammalian proteins. See also Suppl. Table 2.

Of note, the LIDL box we describe here as new clathrin binding motif for drebrin is also present in a variety of other mammalian proteins (Fig. 7b; Suppl. Table 3). Respective AF3 renderings of the AP2-associated kinase AAK1, a risk factor for Parkinsońs disease ^68^ (Ext. Data Fig. 6d-i), or the cell cycle protein GTSE1 (Ext. Data Fig.6j-o) show that, comparable to drebrin, this motif binds to the site of the β-arrestin box on clathrin heavy chain, indicating that the LIDL box is likely of more general importance for clathrin-mediated endocytosis.

## Discussion

In this study, we investigate the molecular machinery that allows direct contact between actin-based podosomes and microtubule +tips. At the same time, we address a broader question, of how endocytosis, selectivity of β integrin uptake, and intracellular trafficking are coordinated at the cell cortex. For this, podosomes present an accessible system, as they form at discrete subcellular sites, whose spatiotemporal relationship with microtubule +tips and the endocytic machinery can be easily visualized and, owing to the high number of podosomes per cell (>300 in macrophages), enables statistical analysis.

Contact by microtubule +tips impacts on both podosome dynamics and function ^29^, comparable to microtubule-dependent regulation of focal adhesions ^1^. However, the dynamic nature of these contacts and the complexity of the subsequent analysis has greatly hindered the identification of factors that differentially regulate podosome-microtubule contact. To address this question, we generated a software tool (“ContactAnalyzer”) that enables the analysis of all podosomes per cell, together with contacting microtubule +tips. ContactAnalyzer is coordinate-based and uses the NET 6 technology that provides console application. ContactAnalyzer solves shortcomings of previous attempts ^31,35^ concerning the detection of multiple simultaneous contacts between podosomes and +tips, allows the analysis of consecutive contacts, enables gathering of all relevant coordinates of podosomes and +tips in all frames of live cell videos, and also enables faster analysis of large data sets. A current limitation of the tool is the need for manual thresholding of the detection radius. Still, the use of ContactAnalyzer is not restricted to the analysis of contact events between podosomes and microtubules +tips but should also be useful for the analysis of contact events between other adhesions such as focal adhesions and other contacting structures such as vesicles.

Importantly, we identify here the actin associated protein drebrin as a new component of the podosome cap. The podosome cap surrounds the core of branched F-actin as an outer substructure of unbranched actin filaments, together with actin-associated proteins, many of which are involved in actin crosslinking or bundling ^25^. The finding that drebrin, an actin bundling protein, shows a more peripheral localization at the cap than α-actinin, a major actin crosslinker, is also in line with the current architectural model of the podosome cap, which predicts a gradient from crosslinked to bundled actin filaments from the inner to the outer layer of the cap and also from the bottom to the top of a podosome ^25^. Drebrin has been previously identified as a component of postsynaptic podosomes in C2C12 murine myotubes, by a general colocalization with F-actin ^58^, but no specific substructural localization was reported. Moreover, downregulation of drebrin was observed to be accompanied by a disordered array of podosome-associated microtubules ^58^. We now expand on these observations by showing that drebrin at the podosome cap is a crucial part of the machinery linking microtubule +tips to podosomes, through its known binding to the +tip protein EB3 ^49^. Accordingly, siRNA-mediated depletion of drebrin leads to reduced numbers of +tip contacts with podosomes.

Interestingly, we identify INF2, another component of the podosome cap, as a negative regulator of the number and duration of podosome/+tip contacts (Fig. 3e,f). The molecular mechanisms behind this phenomenon are currently unclear, but it appears to be in accordance with the generally inverse impact of drebrin and INF2 on the actin cytoskeleton ^69^. It could also be envisioned that tight bundling of the unbranched actin filaments in the cap by INF2 restricts the accessibility of microtubule +tips for drebrin. In line with this, our previous results showed that depletion of INF2 in macrophages leads to an expansion of podosomes in both diameter and height ^26^. Theoretically, INF2 could also work through its ability to stabilize MTs, by binding MTs directly, and also by inducing expression of the tubulin acetyltransferase TAT1 ^70^. However, we do not detect any changes in the level and pattern of MT-stabilizing posttranslational modifications such as acetylation or tyrosination upon INF2 depletion (Ext. Data Fig.7a-d).

Our identification of a drebrin-EB3 axis in podosome regulation is in good agreement with earlier reports demonstrating in vitro binding of His-tagged drebrin to GST-tagged EB3, but not to GST-tagged EB1 ^49^. The drebrin-EB3 interaction was also confirmed biochemically in lysates of rat cortical neurons ^49^, murine C2C12 myotube lysates ^58^ and of human intestinal Caco-2 cells ^57^, and by using FRET/FLIM in filopodia of rat neurons ^49^. Binding of drebrin to EB3 has been shown to link microtubules to F-actin in neuronal growth cones ^48,49^, which is consistent with our finding that drebrin at the podosome cap connects podosomes to microtubules through binding of +tip-localized EB3 (Fig. 3g-l). The specific interaction between drebrin and EB3, and not EB1, has been further demonstrated by the inhibition of neuritogenesis and neuronal migration through an inhibitory construct of EB3 (EB3M), while a corresponding construct of EB1 (EB1M), showed no such effect ^49,71,72^. This specificity for EB3 as a drebrin binding partner seems also to be in good agreement with the finding that EB3 is more proximally located than EB1 at microtubule +tips in growth cone filopodia of embryonic cortical neurons ^73^, and is thus ideally placed to link incoming +tips to F-actin structures such as podosomes. KANK1, another candidate protein included in our screen, was previously shown to link focal adhesions and microtubule +tips, by binding both talin at FAs and liprins at cortical adhesion complexes, with the latter connecting to CLASP at microtubule +tips ^37,38^. Indeed, a previous study has highlighted the importance of KANK1 in linking microtubule +tips to podosomes of THP1 human monocytic cells, with KANK1 depletion leading to strong reductions in podosome numbers and microtubule capturing ^2^. In line with these results, we see a significant decrease in podosome lifetime (Fig. 3a) and density (Ext. Data Fig. 4d) in primary human macrophages upon KANK1 depletion. We also notice a comparable, although non-significant, trend towards reduction of podosome number (Ext. Data Fig. 4c) and podosome/+tip contacts (Fig.3e). Differences between this previous and our current study could be based on the choice of different cell systems, i.e. a monocytic cell line versus primary cells. It is also not clear whether THP1 cells have a drebrin-containing cap structure, although the presence of fascin at a cap-like localization seems to indicate this ^74^. A difference could also be based on our analysis of EB3-positive +tips, which enable contact with podosome-localized drebrin, and whose targeting to podosomes is thus strongly reduced in drebrin-depleted cells.

We further show that depletion of drebrin leads to a general reduced endocytosis of biotinylated surface proteins as well as a specific decrease of β2-integrin endocytosis but not β1 or β3. These results point to a specific role of the drebrin-EB3 axis in β2 integrin endocytosis. At the same time, our results provide novel evidence for the currently unclear basis for integrin isoform preference during clathrin-mediated endocytosis ^75,76^. Of note, clathrin adaptors such as DAB adaptor protein 2 (Dab2) and low-density lipoproteins receptor adaptor protein 1 (LDLRAP1, a.k.a ARH) can interact with the cytoplasmic tail of all β integrins ^75^, thus providing no specificity during integrin uptake. In contrast, drebrin emerges as a selective hub that links endocytosis of β2 integrin at actin structures such as podosomes with microtubule-based transport. Confocal and TIRF microscopy analysis showed a prominent localization of β2 integrin at the ring structure of podosome, a substructure in close contact with both the plasma membrane and the drebrin-positive podosome cap. Similarly, β3 integrin also localizes at the podosome ring, but mostly restricted to larger podosomes at the cell cortex. In contrast, β1 integrin does not show any specific podosomal recruitment. The apparent specificity of drebrin-dependent endocytosis for β2 integrin could thus be, at least in part, based on a close spatial relationship between both proteins.

A recent paper reported that drebrin induces endocytosis of β1 integrin in A549 lung adenocarcinoma cells, through direct or indirect association of both dynamin and β1 integrin with the drebrin C-terminus ^60^. While our current study confirms that the drebrin C-terminus is important for endocytosis, we show in detail that interaction with the clathrin machinery is enabled not by binding to dynamin, but through two clathrin boxes that bind clathrin heavy chain. Indeed, several lines of evidence in our study pointed to a connection between the drebrin-EB3 axis and clathrin-based endocytic events at podosomes: i) Proximity ligation assays showed that drebrin and EB3 (Fig. 3g-n), and also drebrin and clathrin heavy chain (CLH1) (Fig. 5k-q) are in close association at podosomes. ii) Confocal analysis revealed that EEA1-positive endosomes are often associated with podosomes, and that CLH1 and EEA1 show significant colocalization, especially at the drebrin-containing podosome cap (Fig. 5r-w). iii) Live cell imaging demonstrated that clathrin vesicles are often in close contact with podosomes and emerge adjacent to the drebrin-positive podosome cap (Fig. 5x). iv) Co-immunoprecipitation of drebrin-GFP from macrophage lysates indicated an association of drebrin with the clathrin machinery (Fig. 4j). This was further substantiated by AlphaFold3-based modelling, which pointed to potential binding of two motifs in the drebrin C-terminus to hydrophobic pockets in the clathrin heavy chain, both with high ipTM scores (Fig. 6a,b). vi) Mutation of these motifs, exchanging critical amino acid residues with glycine or alanine residues, led to a reduction in coprecipitation of CLH1 to background levels (Fig. 6e,f).

Interestingly, the CLH1-binding boxes in the drebrin C-terminus share sequence motifs with several of the previously defined clathrin-binding boxes. Specifically, the sequence ^530^ LLNFDE^535^ matches the “canonical box” motif LΦXΦ[DE] and is predicted by AF3 to bind to the hydrophobic pocket of the β-propeller 1 and 2 of the CLH1 N-terminus. Interestingly, the second clathrin binding sequence ^475^LIDLWP^480^ matches partially with the “canonical box” motif LΦXΦ[DE], without the last negatively charged amino acid [DE], with the “W-box” motif PWXXW, without the second tryptophan [W], and with the “β-arrestin box” [LI][LI]GXL, without the glycine [G]. Moreover, AF3 modelling of the drebrin C-terminus (421-540) with the CLH1 N-terminus (1-360) predicts an interaction of drebrin ^475^LIDLWP^480^ with the “β-arrestin box”, between the β-propeller 4 and 5 of clathrin, indicating that both drebrin boxes can bind to different motifs on the CLH1 surface, predicting a certain degree of promiscuity between the drebrin boxes, especially concerning the ^475^LIDLWP^480^ sequence. This seems to be in line with a previous report demonstrating that binding of the four classical clathrin boxes to the CLH1 surface is degenerate and depends on the structural context in which the respective peptides are included^77^.

Of note, our discovery of the LIDL box in drebrin prompted us to search for similar motifs also in other mammalian proteins. Indeed, a position-specific scoring matrix (PSSM) search ^78^ returned a list of 47 proteins annotated to interact with clathrin, although with unknown interaction sites, and containing the motif [LMIV][IFVM]D[LFIM] (Fig. 7b, Suppl. Table 3). Interestingly, this motif is similar to the canonical box motif LΦXΦ[DE], but with the negatively charged amino acid situated in the middle rather than at the end of the motif. The shift in the position of D or E matches the different configuration of positively charged amino acids close to the hydrophobic pockets on the clathrin surface. Specifically, the R^64^ and K^96^ of the β-propeller 1 and 2, responsible of the electrostatic interactions with the negatively charged amino acids [DE] of the canonical box are situated on the side of the hydrophobic pocket, whereas the R^188^ and K^245^ of the β-propeller 4 and 5 (β-arrestin box) are situated in the middle of the hydrophobic pocket (Ext. Data Fig. 6a,b).

Proof-of principle analyses using AF3 showed that the respective boxes of such diverse proteins as the AP2-associated kinase AAK1, a regulator of clathrin-mediated endocytosis ^79^, or G2- and S-phase expressed protein (GTSE)1, are modelled to the site of the β-arrestin box, similar to drebrin (Ext. Data Fig.7d-o). These results indicate a more widespread importance of this box for clathrin-mediated endocytosis in a variety of scenarios, also considering that AAK1 is involved in virus entry into host cells ^80,81^, and that GTSE1, a regulator of p53 activity ^82^, is a biomarker for various tumors ^83^. Moreover, our results are in line with a recent report that identified the LIDL motif in a variety of proteins from the *Saccharomyces cerevisiae* genome, and confirmed binding of the LIDL box to the site of the β-arrestin box on CLH1 ^84^.

These results appear to be in good agreement with a report showing that branched actin networks, which can also be found in podosome cores, can be associated with clathrin-coated structures and can promote invagination of flat clathrin arrays in mammalian cells ^85^. Of note, this model would not necessitate drebrin binding to both clathrin and EB3 at all stages of the endocytic process. It could be envisioned that drebrin first binds CLH1 near the plasma membrane, as endocytic vesicles form and pinch off. These vesicles could then stay in dynamic contact with the podosome cap through consecutively binding drebrin molecules at more cell-internal levels of the cap, until reaching a drebrin population that is bound to EB3, thus enabling their transfer to microtubules. However, further modelling using AlphaFold3 points to an alternative scenario: The drebrin C-terminus, containing the two clathrin binding sites, is only exposed when drebrin is bound to an actin trimer, but not to a filament of at least four actin subunits. Accordingly, it could be reasoned that CLH1 could only bind to drebrin that is associated with actin nuclei, i.e. during phases of actin filament formation or depolymerisation (Ext. Data Fig.9, Suppl. Video 7). This model would be in line with the observation that clathrin is found in close proximity to podosomes especially during the initial stages of their formation, when actin nuclei are likely more abundant, or at the cytoplasm-facing surface of the podosome cap, where actin depolymerisation occurs due to aging of the filaments and distance from actin nucleators ^25^ (Fig. 5x_2_, Suppl. Video 4).

Moreover, not only drebrin, but also other proteins of the podosome cap could contain clathrin boxes, as has been demonstrated for α-actinin ^86^. Drebrin might thus not be the only cap protein mediating clathrin binding at podosomes. Still, drebrińs dual ability to also bind EB3 makes it a critical hub that links clathrin-based endocytosis to +tip recruitment. It should also be noted that the presence of podosomes is not a prerequisite for endocytosis. In addition to its localization at the podosome cap, drebrin at the cell cortex could act as a hub for clathrin-dependent endocytosis and microtubule-based transport in cells that do not form podosomes or in non-podosome containing parts of cells. Still, the local concentration of drebrin at the podosome cap likely facilitates this process.

Our results show that the clathrin-drebrin-EB3 axis works as an on-ramp that enables access of endocytic vesicles to the microtubule system. This raises the question of the potential nature of an off-ramp for exocytic vesicles that are about to leave microtubule-based transport. As mentioned, the end binding proteins EB1 and EB3 are not uniformly distributed on +tips ^73^. Combined with the observation that +tips in primary macrophages show heterogeneous decoration with CLIP170 and EB3 (Ext. Data Fig.7e-k), this could indicate the existence of microtubule subsets with different amounts of specific +tip proteins and likely also different functions. Moreover, considering the relatively large size of microtubules, it could be envisioned that both on-and off-loading of cargo could take place at the same +tips. Indeed, it has been proposed that multiple redundant pathways exist to control cortical targeting of microtubules ^87^. This could also, in part, explain the finding that surface levels of integrin β1 are unchanged upon depletion of drebrin or EB3. We show that β1 integrin, in contrast to β2 integrin, does not associate with drebrin, and thus has no indirect link to EB3. Instead, β1 integrin could be linked to MTs via other adaptors such as EB1, which has been shown to be part of its interactome ^87^.

Interestingly, it has been shown that human immunodeficiency virus (HIV)-1 uses podosomes as entry sites into macrophages ^44^, and that the HIV pathogenicity factor Nef can increase stability and size of macrophage podosomes ^88^. As HIV-1 can enter cells also by clathrin-mediated endocytosis ^89^, this may point to the potential involvement of the clathrin-drebrin-EB3 axis in HIV-1 entry into macrophages and also other podosome-forming cells. This should be an interesting point worth further study.

In conclusion, we show that drebrin is an actin-based, multivalent and selective hub that links clathrin-dependent endocytosis with microtubule-based transport. We identify drebrin as a novel component of the podosome cap that, together with its binding partner EB3, regulates podosome-microtubule +tip contact. Moreover, drebrin binds the clathrin heavy chain by two motifs in its C-terminus and works as a spatially selective regulator of β2 integrin uptake during clathrin-dependent endocytosis, similar to the role of swiprosin during clathrin-independent endocytosis of β1 integrin ^90^. Moreover, our identification of a novel mammalian clathrin box, which we show is present in a wide variety of proteins, should be of general importance for the field of clathrin-mediated endocytosis.

## Online Methods

### Isolation and culture of macrophages

Monocytes were isolated from leukocyte fraction of buffy coats (kindly provided by Frank Bentzien, UKE Transfusion Medicine, Hamburg, Germany), by sucrose density gradient (Ficoll by PromoCell) and isolated using anti-CD14 magnetic beads and separation columns (Miltenyi Biotec), as described previously ^91^. Cells were cultured in monomedium (RPMI-1640 containing 100 units/ml penicillin, 100 μg/ml streptomycin, 2 mM glutamine and 20% autologous serum) at 37 °C, 5% CO_2_ and 90% humidity. Monocytes were differentiated into macrophages by culturing them for at least seven days.

### Transfection

For the depletion of target proteins, macrophages were transiently transfected with respective siRNAs (see Suppl. Table 1) using the Neon electroporation system (ThermoFisher Scientific, MA, USA). Briefly, cells were detached with 1 ml 1X Accutase for 1h at 37°C (Serana Europe GmbH, Pessin, Germany), washed with PBS and counted. 5µl of 20µM siRNA were transfected to 1x10^6^ macrophages using 2 pulses of 40 msec at 1000V. Transfected cells were immediately transferred into 1 ml RPMI and seeded onto a single well of a 6-well plates. After 2-3 h RPMI was replaced by medium containing human serum. Cells were used for subsequent experiments 72-96 h after siRNA transfection.

Experiments employing the overexpression of fluorescently labelled constructs (see Suppl. Table 1) were performed by transfecting 0.5 µg of plasmid to 1x10^5^ cells following the settings and the system described above. Transfected cells were immediately transferred into RPMI and seeded in a µ-Slide 8 Well high (Glass Bottom, Cat# 80807) (Ibidi GmbH, Germany). After 2-3 h, RPMI was replaced by medium containing human serum.

### Cloning

All constructs were generated using a standard molecular cloning approach with kanamycin resistance. The vector was amplified via PCR using 5’-phosphorylated primers and the methylated parental plasmid digested with FastDigest *Dpn*I (ThermoFisher Scientific, Cat# FD1703) at 37°C for 1 hour, while the resulting PCR product was run on a 1% agarose gel, the appropriate band excised and extracted using the QIAquick Gel Extraction Kit (QIAGEN, Cat#28704). Ligation was performed at 16°C overnight using T4 DNA Ligase (Promega, Cat#M180A). The ligation product was then transformed into DH5α *E. coli* strain via heat shock, plated on kanamycin plates, and grown overnight. Colonies were picked for mini prep, inoculated into 2 mL cultures, grown overnight, and plasmids were extracted using the ZR Plasmid Miniprep - Classic (Zymo Research, Cat#D4054). Sequencing confirmed the correct sequences, followed by maxi prep for further experiments and storage. For specific primer sequences, see Suppl. Table 1.

### Protein sample preparation and Western blot

To prove the efficiency of knockdown experiments, siRNA transfected cells were lysed with a lysis buffer (150 mM NaCl, 1% Triton X-100, 50 mM Tris-HCl (pH 8.0), containing protease and phosphatase inhibitors (Roche, Germany)). After incubation on ice for 10min, cells were scraped and collected in tubes. Cell debris were then pelleted by centrifugation at 14,000 g for 30 min at 4°C, supernatants were mixed with sample loading buffer (5x) and heated for 10min at 96°C for denaturation. Proteins were then separated by SDS-PAGE using custom 10% acrylamide gels, following the manufacturer’s protocol (Bio-Rad Laboratories Inc, California, USA) or 4-12% Bis-Tris precast gradient gels (Merck Millipore, Cat# MP41G15) and then transferred onto nitrocellulose or PVDF membranes using the iBlot2 dry blotting system (ThermoFisher Scientific, USA). Membranes were blocked using 1X BlueBlock (Serva Electrophoresis GmbH, Heidelberg, Germany) or 5% low-fat dry milk (Carl Roth, Cat# T145.2) for at least 3 hrs at RT and then incubated with primary antibodies diluted in 1x BlueBlock or 5% BSA (Sigma Aldrich, Cat# A9647) at 4°C overnight, while HRP-coupled secondary antibodies were probed for at least 1h at room temperature. Between primary and secondary antibody incubation, membranes were washed at least three times with TBS-T for at least 20 min. Western blots were then acquired using an Amersham ImageQuant™ 800 imaging system (Cytiva, Sweden) after incubation with Pico or Femto chemiluminescence solutions-(ThermoFisher Scientific, USA). When needed, nitrocellulose membranes were mild-stripped by extensive washing with stripping buffer (200 mM glycine, 3.5 mM SDS, 1% tween20, (pH 2.2)) before reblocking and reprobing, whereas PVDF membranes were stripped with a guanidine hydrochloride-based (GnHCl) stripping solution (6M GnHCl, 0.3% Triton X-100, 0.1M DTT (added fresh), 20mM Tris-HCl, pH7.5) cite this paper (doi:10.1016/j.ab.2009.03.017).

### Immunoprecipitation and GFP-IP

Immunoprecipitation of endogenous protein was performed according to manufacturer’s instructions (Miltenyi Biotech), with modifications. Two-weeks old cells from three six-well plates were washed three times in prewarmed PBS, lysed with Triton X-100 lysis buffer [150 mM NaCl, 1% Triton X-100, 50 mM Tris-HCl (pH 8.0)] containing protease and phosphatase inhibitors (Roche) (300 µL / well) and shaken on ice for 15 min before being scraped and collected in tubes. The sample was sonicated twice for 10 sec at 25% amplitude, rotated at 4 °C for 15 min before saving a small aliquot as “input”, and then precleared with a mixture of 200 µL µMACS Protein A/G magnetic microbeads in constant rotation for 30 min at 4 °C. A µMACS column was prewashed with lysis buffer, loaded with the protein sample and the eluate collected. The cleared sample was then split into three equal volumes and incubated with a mixture of 200 µL µMACS Protein A/G magnetic microbeads + 10 µg of Normal Rabbit IgG (Cell Signalling) as Isotype control, or 10 µg of anti-Drebrin Ab (Proteintech) or 10 µg of anti Dynamin 2 Ab (Abcam). Mixtures were then rotated overnight at 4°C. After incubation samples were loaded into prewashed µMACS columns, washed once with lysis buffer, twice with RIPA buffer [150 mM NaCl, 1% Triton X-100, 0.5% sodium deoxycholate, 0.1% SDS, 50 mM Tris-HCl (pH 8.0)] and once with 20 mM Tris-HCl (pH 7.5) before elution with the buffer from the kit [50 mM Tris–HCl (pH 6.8), 50 mM DTT, 1% SDS, 0.005% bromophenol blue, 10% glycerol] pre-heated at 95°C. Eluted samples were then mixed with 4× Laemmli sample loading buffer, heated 10 min at 95 °C and examined by Western blot.

Immunoprecipitation of GFP-fused proteins was performed according to manufacturer’s instructions (Miltenyi Biotech), with modifications. Cells overexpressing overnight drebrin-GFP or, as control, pEGFP empty plasmids, were washed three times in prewarmed PBS, lysed with Triton X-100 lysis buffer [150 mM NaCl, 1% Triton X-100, 50 mM Tris-HCl (pH 8.0)] containing protease and phosphatase inhibitors (Roche) and shaken on ice for 15 min before being scraped and collected in tubes. Samples were then rotated for 90 min at 4°C, centrifuged at 10.000 RPM for 10 min at 4°C to pellet cell debris, supernatants saved and incubated with µMACS anti-GFP microbeads, for 2 h in constant rotation at 4 °C. After incubation, samples were loaded into prewashed µMACS columns, washed once with lysis buffer, twice with regular (“mild”) or high salt (“stringent”) RIPA buffer [500 mM NaCl, 1% Triton X-100, 0.5% sodium deoxycholate, 0.1% SDS, 50 mM Tris-HCl (pH 8.0)] and once with 20 mM Tris-HCl (pH 7.5) before elution with buffer [50 mM Tris–HCl (pH 6.8), 50 mM DTT, 1% SDS, 0.005% bromphenol blue, 10% glycerol] pre-heated at 95°C. Eluted samples were then mixed with 4× Laemmli sample loading buffer, heated 10 min at 95 °C and examined by Western blot.

### Surface biotinylation and endocytosis assays

Endocytosis assay, analysed by Western blot, was performed using reagents from Pierce Cell Surface Protein Biotinylation and Isolation Kit (ThermoFisher Scientific, Cat# A44390), with modifications. Macrophages, transfected with siRNA 3 days prior to experiments, were put on ice for 15 min to stop endocytosis, then washed twice with ice-cold DPBS containing calcium and magnesium (DPBS++) (ThermoFisher Scientific, Cat# 14040133) and incubated for 30 min on ice with 1 mM cleavable EZ-Link Sulfo-NHS-SS-Biotin, dissolved in DPBS ++, to covalently biotinylate surface exposed proteins. After incubation, cells were washed once with ice-cold DPBS++ before quenching the free Sulfo-NHS-SS-Biotin with ice-cold TBS buffer washes (2x5 min). After two additional washes with ice-cold DPBS++, cells were lysed and the positive control prepared for downstream application, whereas the negative control, to check the cleavage efficiency, was further incubated 3 ×10 min on ice with ice-cold 100 mM MESNA dissolved in TBS buffer (50 mM Tris-HCl, 100 mM NaCl, 2.5 mM CaCl2, pH 7.8), to reduce S-S bonds and strip biotin from biotinylated proteins on the cell surface. After three additional washes with ice-cold DPBS++, cells of the negative control were lysed and samples collected for downstream applications. To analyse specific time points of endocytosis, following the quenching step and the two washes with ice-cold DPBS++, biotinylated cells were incubated with prewarmed standard monomedium at 37°C for the indicated times. Cells were then put on ice to stop endocytosis and washed once with ice-cold DPBS++, followed by 3 ×10 min incubation with ice-cold MESNA to strip biotin from non-internalized proteins. After three additional washes with ice-cold DPBS++, cells were lysed and samples collected for downstream applications. All samples were lysed for 5 min on ice using a 1:1 mixture of ice-cold lysis buffer (from kit) and RIPA buffer, containing protease inhibitor cocktail (Cat# 04693124001, Roche). Lysates were then scraped, collected in tubes and incubated on ice for 30 min, with 5 sec vortexing before and after the incubation. Cell extracts were then centrifuged at 15,000 g for 5 min at 4°C before transferring supernatants into new tubes. Protein concentration of lysates was measured using Pierce BCA assay (ThermoFisher Scientific, Cat# 23225), and samples with equal amounts of protein were prepared and incubated overnight at 4°C in columns together with NeutrAvidin Agarose resin, with end-over-end mixing. The day after, samples were washed four times with wash buffer (from kit) and eluted with elution buffer (from kit) containing 10 mM of DTT, to allow the reduction of S-S bonds and the release of proteins from the EZ-Link Sulfo-NHS-SS-Biotin. Eluted samples were mixed with sample loading buffer (5x) and heated 10 min at 95°C prior to Western blot analysis. Endocytosis rates were calculated by comparison to the positive control, set as 100%, which represents the surface amount of a given protein at time point 0.

Endocytosis assays, analysed by immunofluorescence, were carried out as described above, but with the following modifications. After siRNA transfection, cells were seeded on several µ-Slides 8 Well high (Glass Bottom, Cat# 80807) (Ibidi GmbH, Germany). At due time points, samples were fixed for 10 min at room temperature with 4% methanol-free formaldehyde, to avoid undesired cell permeabilization at this stage. Following fixation, samples were washed three times with DPBS, regularly permeabilized for 5 min with 0.5 % Triton X-100, blocked for 1 hr with 2% BSA and 5% NGS, incubated for 90 min with blocking solution containing 1:400 AlexaFluor488-streptavidin (ThermoFisher Scientific, Cat# S32354), 1:400 AlexaFluor568-phalloidin (ThermoFisher Scientific, Cat# A12380), 1:1000 DAPI (ThermoFisher Scientific, Cat# 62248) and 1:200 anti-drebrin polyclonal Ab (ThermoFisher Scientific, Cat# PA5-84067), washed with 0.05% Triton X-100 (3x5 min) and then incubated for 60 min with blocking solution containing AlexaFluor647-anti-rabbit Ab (ThermoFisher Scientific, Cat# A-21246) before 3x5 min washes with 0.05% Triton X and mounting with Fluoromount-G medium (ThermoFisher Scientific, Cat# 00-4958-02). Z-stacks of Immunofluorescent samples were then acquired at Visitron Spinning disk with a 40x oil objective and with a Z-step of 0.25 µm.

### Proximity ligation assay

Proximity labeling was performed following the manufactureŕs instructions (Duolink, Sigma-Aldrich, Darmstadt, Germany). Following fixation with 4% paraformaldehyde (PFA), cells were permeabilized with 0.5% Triton X-100 and blocked using Duolink blocking buffer. Primary antibodies used included MAB3580 (Chemicon, mouse) against GFP, ab157217 (Abcam, rabbit) against EB3, MA1-065 (Invitrogen, mouse) against clathrin heavy chain and PA5-84067 (Invitrogen, rabbit) against drebrin. After primary antibody incubation, Duolink In Situ Detection Reagents Red (MilliporeSigma, DUO92008) were applied according to the manufacturer’s protocol. Secondary probes specific for mouse (‘-‘ probe) and rabbit (‘+’ probe) antibodies were introduced. During the probing phase, Alexa Fluor 405-labeld phalloidin was added at a dilution of 1:400 to stain actin filaments. Samples were mounted using Fluoromount-G Mounting Medium (ThermoFisher Scientific, Cat# 00-4958-02) for imaging. To assess the spatial association between Proximity Ligation Assay (PLA) signals and podosomes, we used a custom-developed command-line image analysis tool available via GitHub (“BioPixel”; https://github.com/bryanbarcelona/BioPixel). BioPixel enables detection and spatial mapping of PLA puncta relative to podosome structures through an interactive command-line interface. While not yet featuring a graphical user interface, it supports reproducible and flexible workflows tailored for fluorescence microscopy image analysis.

### Immunofluorescence and microscopy

Samples were fixed for 10 min in 3.7% formaldehyde, with a 3 sec prefixation in ice-cold methanol. Coverslips were mounted using Mowiol (Calbiochem, Darmstadt, Germany) containing 1,4-diazabicyclo[2.2.2]octane (25 mg/ml; Sigma-Aldrich, Saint Louis, MO, USA) as anti-fading reagent or Fluoromount-G medium (ThermoFisher Scientific, Cat# 00-4958-02). Images of fixed samples were captured using a Leica DMi8 confocal laser-scanning microscope, equipped with a TCS SP8 AOBS confocal point scanner, an oil-immersion 63× HC PL APO Oil CS2 NA 1.40 objective, and both HyD and PMT detectors. Imaging was performed using Leica LAS X SP8 software (version 3.5.2, Leica Microsystems, Wetzlar, Germany). AiryScan super-resolution images of fixed samples were acquired using a Zeiss Axio Examiner Z1 with LSM 980 and Airyscan 2 (upright), equipped with a 63x Plan-APOCHROMAT Oil DIC objective (NA: 1.40), Airyscan 2 detectors and the Zeiss ZEN software.

For live cell imaging, cells were transfected with respective constructs, as indicated, seeded on a µ-Slide 8 Well high, Glass Bottom (Ibidi GmbH, Cat# 80807) at subconfluent concentration and incubated overnight before acquisition. Time lapse movies were acquired using the Visitron spinning-disk system equipped with a Nikon Eclipse TiE, a 100x CFI Plan Apo Lambda objective (NA: 1.45) a Yokogawa CSU W-1 SoRa for super-resolution (pixel size: 0.039), 2x Photometrics Prime 95B (back-illuminated sCMOS, 11µm pixel-size, 1200x1200 pixels) cameras, an environmental chamber with temperature, humidity and CO_2_ control, and the VisiView v4 software. Staining of integrins was performed by fixation with 4% methanol-free formaldehyde for 15 min, permeabilization with 0.5% Triton X-100 for 5 min, blocking with 2% BSA together with 5% NGS (Natural Goat Serum) for 1 h and incubation with AlexaFluor647-conjugated primary antibodies together with AlexaFluor488-conjugated phalloidin also for 1 h. TIRF imaging was performed with an iLAS TIRF unit from Visitron systems fitted on a Nikon eclipse Ti microscope with oil immersion Plan-Apo 63× NA 1.45 and Plan-Apo 100× NA 1.49. For analysis of podosome/microtubule +tip contacts, cells were transfected to overexpress LifeAct-TagRFP (Ibidi GmbH, Cat# 60102) together with EB3-GFP, CLIP170-GFP, or EB1-GFP and seeded in µ-Slides 8 Well high, Glass Bottom (Ibidi GmbH, Cat# 80807). After 4.5 hrs of incubation, acquisition was started and 30 min videos were taken with a frame rate of two images every 2 seconds, with respective TIRF imaging at 488nm and 568nm.

### Image analysis and quantification

To analyse the three-dimensional distribution of GFP-fused drebrin constructs at podosomes, the ImageJ-based Poji macro was used ^53^. Briefly, in the focal plane of highest F-actin intensity, the macro analyzes the positions of individual podosomes by detecting their respective brightest pixel (“Find Maxima” tool). It measures the fluorescence intensity profile across a line of 15 pixels (corresponding to ∼1.65 µm), rotating it 360 times for 1 degree each and repeating the measurement after every rotation and every focal plane. The mean of all 360 measurements per focal plane is then averaged across all acquired cells and plotted in a stacked graph of 5 z-planes, with a Z-step of 0.20 µm (∼ 1 µm in height with 0.5 µm spanning towards dorsal and ventral direction, respectively). The focal plane with the highest F-actin intensity is labelled as z=0, also for the corresponding GFP channel, with the focal planes in dorsal and ventral position being marked as z=+x and z=-x, respectively. To avoid fixation artifacts, the Z-stacks of cells overexpressing single GFP-constructs together with LifeAct-TagRFP (Ibidi GmbH, Cat# 60102) were acquired at Visitron Spinning Disk with a 100X objective, in live-imaging settings and dual camera simultaneous acquisition mode.

To analyse and quantify endocytosis by immunostaining, Z-stacks of whole cells from all channels (DAPI, AlexaFluor488-streptavidin, AlexaFluor568-phalloidin and endogenous drebrin) were projected onto a single Z-plane by averaging intensities. Images were then uniformly contrasted by setting fixed minimum and maximum intensity values. Based on the F-actin signal (AlexaFluor568-phalloidin) single cell ROIs were manually drawn and recorded, whereas the nuclei signal (DAPI) was used as a reference to include only intact cells. Next, the mean intensity values of both drebrin and streptavidin channel were imported into GraphPad for each single ROI. Considering that the intensity values of a 16-bit image can range from 0 to 65535, a middle value of 30000 was set as a threshold to prune control cells (> 30000) in the control sample and drebrin knockdown cells (<30000) in the knockdown sample. Endocytosis rates were calculated by comparison to the respective control, set as 100%, which represents the whole amount of surface proteins at time point 0.

### Podosome- +tip contact analysis using the ContactAnalyzer tool

The ContactAnalyzer tool was created for contact analysis between podosomes and microtubule +tips, from live cell videos. This tool uses spots (coordinates) and tracks (spots from individual frames that are combined over time), created by the ImageJ plugin TrackMate in XML format ^92^. ContactAnalyzer processes one reference file (podosome channel) together with one channel file (+tip channel). Overlap of two spots is calculated via a circular intersection of both spot coordinates and their given detection diameter. For calculating a raw contact, a podosome track is selected. Subsequently, each spot of this track is checked for potential overlaps with all +tip spots of the respective frames (Ext. Data Fig.1b). The diameter (in pixels) for the overlap calculation is adjustable and depends on the zoom and resolution of the movies to analyze. For the above-described imaging settings, detection diameters of 12 pixels for podosomes and 5 pixels for the +tips were chosen, matching the visual confirmation of a contact. If the spots of podosomes and +tips have an intersection area of >0, this is identified as a raw contact, and respective coordinates are stored. The program continues this analysis frame by frame until the end of the track. Subsequently, this process will be repeated for the next track until all tracks are examined. For reasons of performance speed, this process runs parallel for several tracks. After the detection of all raw contacts (i.e. all +tip spots that have an overlap with a podosome), they are checked for being potentially consecutive. A contact is consecutive if a spot from a track in frame n also is in contact with the spot in frame n+1 for any number of following raw contacts in subsequent frames without interruption, thus the corresponding time equals the contact duration. Consecutive contacts will be calculated with the overlap method as well and stored as an intersection matrix of all contacts (Ext. Data Fig.1e,f). With iterations selecting the highest value in the matrix and removal of processed values, respective raw contacts can only be part of a single consecutive contact. For the ContactAnalyzer code, see https://github.com/mbwcode/ContactAnalyzer.

### SLiM Discovery

Bioinformatic analysis to find novel linear motifs was performed as previously described ^84^, with modifications. Position-specific scoring matrix (PSSM) search ^78^ was used to search for proteins containing the LIDL motif within the *Homo sapiens* proteome. Specifically, the LIDL peptide was entered as input selecting in the "output advanced options" 5 amino acids as flank length and adding the UniProt accession number of clathrin heavy chain (Q00610) as shared annotations with a P-value cut-off of 0.05, to filter out proteins not annotated to interact with clathrin heavy chain. Successively, the output list was cleaned online by activating filters for "Warning" (disorder, domain, topology, surface accessibility, localisation) and "Hub" (interaction, localisation, function, process and any ontology). The list with 95 hits in 72 proteins was then imported into an excel datasheet and manual curation was performed for obtaining the final protein list, discarding proteins where the motif is in a structured region or those where the motif matches an already described clathrin-binding motif, such as the canonical box, the W-box, the β-arrestin box or the DLL motif.

### Protein structure prediction and visualization

3D models of Drebrin (DBN1) (Uniprot Q16643, aa 421-540) in complex with Clathrin heavy chain 1 (CLTC) (Uniprot Q00610, aa 1-360) were generated using AlphaFold 3 ^93^ via the AlphaFold server (https://alphafoldserver.com) to capture accurate protein-protein interactions. The overall accuracy of the model structure and of the predicted interaction between chains were based on pTM (Predicted Template Modeling) and ipTM (Interface Predicted Template Modeling) scores, respectively. Specifically, the mentioned model had an ipTM of 0.72 and a pTM of 0.76, confirming a good quality of the predicted interfaces. The model of AP2-associated protein kinase 1 (AAK1) (Uniprot Q2M2I8, aa 823-961) in complex with Clathrin heavy chain 1 (aa 1-360) had an ipTM of 0.68, and a pTM of 0.75, whereas G2 and S phase-expressed protein1 (GTSE1) (Uniprot Q9NYZ3, aa 1-739) also in complex with Clathrin heavy chain 1 (aa 1-360) had an ipTM of 0.73, and a pTM of 0.42, the latter most likely due to the highly unstructured nature of GTSE1.

Molecular visualization and interaction analysis were conducted in UCSF ChimeraX ^94^. Hydrophobic interactions and van der Waals contacts were mapped using the “contacts” command, whereas electrostatic potential surfaces were generated to identify potential charge-based complementarity at the binding interfaces.

### Statistics

Datasets were tested for normal (Gaussian) distribution by plotting the frequency distribution and performing a normal/log-normal Quantile-Quantile-Plot (QQ plot). Most datasets showed a lognormal distribution, as commonly observed in biological data. Therefore, comparing more than two conditions, the non-parametric Kruskal-Wallis test was performed, followed by Dunn’s multiple comparison post hoc test. Natural logarithm transformation was performed using an ordinary one-way ANOVA and a Dunnett’s multiple comparison post hoc test, with both tests leading to comparable significances. Note that the second variant is presented only in the supplement, owing to the reduced readability of non-linear scales. For endocytosis assays, a 2way ordinary ANOVA was performed with the Sidak post-hoc test for multiple comparisons. The endocytosis assay analysed by Western blot includes data from 2 different donors (n=2), whereas the endocytosis assay analysed by immunofluorescence includes 50 cells per time point and per condition (control vs siRNA-induced knockdown) from 4 different donors (n=200 per time point and condition). For the statistical analysis of the GFP-IP of Drebrin-PP + Cterm GFP wildtype and mutants, a One sample t and Wilcoxon test was performed on quantifications collected from three different donors (n=3). All data were collected using GraphPad Prism or Microsoft Excel, while statistical analysis was performed using GraphPad Prism. For details, see Suppl. Table 2.

## Supporting information

Ext Data Fig 1-9

## Acknowledgements

We thank Frank Bentzien (UKE transfusion medicine) for buffy coats, Niels Galjart for CLIP170-GFP and EB3-mRFP, Marina Mikhaylova for EB3-GFP, Yuko Mimori-Kiyosue for EB1-GFP, Kostiantyn Sopelniak for help with PLA, Andrea Mordhorst and Kevin Schiemang for expert technical assistance, the UKE microscopy facility (umif) for help with microscopy and image analysis, Michael Windler for help with software analysis, and Martin Aepfelbacher for continuous support. Microscopes used in this study within the umif were available thanks to support by Deutsche Forschungsgemeinschaft: INST 337/34-1 FUGG: Leica TCS SP8 X, INST 337/20-1: Visitron SD-TIRF, INST 152/876-1 FUGG: AiryScan).

Molecular graphics and analyses performed with UCSF Chimera, developed by the Resource for Biocomputing, Visualization, and Informatics at the University of California, San Francisco, with support from NIH P41-GM103311. This work is part of the doctoral theses of K.W. and B.B. and was supported by Deutsche Forschungsgemeinschaft (LI925/8-1; LI925/13-1).

## Author contributions

P.C., S.H., K.W., and B.B. performed experiments, R.H. performed data analysis, P.C. and S.L. devised concepts and experiments, S.L. and P.C. wrote the manuscript.

## Extended data

**Extended Data Figure 1. Podosome/+tip contact analysis using the ContactAnalyzer tool.** (a-d) Required features for a program detecting podosome/+tip contacts. Red pixels: podosome detection via lifeact-RFP, green pixels: MT +tip detection via EGFP signals of respective fusion proteins. Red and green circles indicate respective detection radii, (a) detection of +tip contact at a given time point n; (b) detection of multiple +tip contacts at a given time point n; (c) identification of a persistent contact by the same +tip, based on identical localization in subsequent frames at time points n and n+1; (d) identification of contacts by different +tips, based on different localization in subsequent frames at time points n and n+1; (e) Contact detection by the ContactAnalyzer tool: Two subsequent frames with a detection radius of podosome track with two spots (S1 in frame n, S10 in frame n+1) and several MT +tips (Spot +Tip). S2, S3, S13 and S14 show an overlap with spots of the podosome track. Four raw contacts (S1, S2); (S1, S3); (S10, S13) and (S10, S14) will be stored as raw contact. Using matrix calculation, the MT +tip spots S3 and S13 will be recognized as the same spot and therefore saved as a consecutive contact. S2 and S14 will be recognized as two different contacts. (f) Matrix calculation: An example of three iterations of the used matrix calculation is depicted. A frame n contains four raw contacts (S1-S4) and the subsequent frame n+1 contains three raw contacts (S5-S7). The intersection area of all possible combinations between the spots from frame n with those of frame n+1 is calculated as described in https://github.com/mbwcode/ContactAnalyzer. With iterations selecting the highest value in the matrix and removal of processed values, respective raw contacts can only be part of a single consecutive contact.

**Extended Data Figure 2. Evaluation of +tip proteins for analysis of podosome/+tip contact.** (a-c) Still images of TIRF movies of macrophages expressing CLIP170-GFP (a), EB1-GFP (b), or EB3-GFP (c). Scale bar: 10 µm. (d-i) Analysis of cellular and podosomal parameters in cells expressing the indicated constructs imaged in TIRF for 30 min (1 fps). (d) podosome numbers, (e) number of observed +tips, (f) number of raw contacts, (g) number of consecutive contacts, (h) consecutive contacts per podosome, (i) mean contact duration. Values were analysed using the non-parametric Kruskal-Wallis test, followed by Dunn’s multiple comparison post hoc test, *P<0.05, **P<0.01, ****P<0.001. For specific values, see Suppl. Table 2.

**Extended Data Figure 3: Domains involved in targeting drebrin to podosomes.** Confocal micrographs of macrophages overexpressing respective drebrin-GFP deletion constructs, as indicated, and Lifeact-RFP to label podosome cores; white boxes in overview images indicate detail regions shown on the right; scale bars: 6 µm for overviews, 1 µm for detail images. (a) Drebrin contains an N-terminal ADF homology region (ADFH; aa 1-135), a coiled coil (CC) region regulating homodimerization (aa 176-256), a helical domain (Hel; aa 257-355), a polyproline region (PP; aa 364-417) and a long unstructured C-terminal region (431-649). Tested constructs included (b) a deletion of the ADFH domain (ΔADFH-GFP), (c) a deletion of both ADFH and PP domains (ΔADFH-ΔPP-GFP), (d) a construct containing only the coiled coiled and helical domains (CC-Hel-GFP), (e) a construct containing the PP and C-terminal regions (PP-C-term-GFP), and (f) a construct containing only the C-terminus of drebrin (Cterm-GFP); domain cartoons of respective constructs are shown on the side. Graphs show fluorescence intensity of GFP-fusion proteins at podosomes, as analysed using Poji macro in several optical planes with a distance of 250 nm each. Plane with maximal intensity for F-actin is marked as z=0, with colour code for optical planes depicted on the right of (b) (> 200 podosomes from at least 3 cells). Note wing-shaped profiles in (b,c), indicating localization at the podosome cap, dome-shaped localization in (d), indicating localization at the podosome core, and diffuse localization in (e,f), indicating more unspecific localization.

**Extended Data Figure 4. siRNA-mediated depletion of candidate proteins; additional evaluated parameters.** (a) Western blots of lysates from macrophages treated with indicated siRNAs, or control siRNA; remaining protein levels after a 3d knockdown are given on the right. (b-e) Evaluation of podosome/+tip contacts during 30 min TIRF videos, using the ContactAnalyzer tool. (b) cell area in µm², (c) podosome number, (d) podosome density (podosome number per cell area in µm²) and (e) number of microtubule +tips per cell. Statistics: non-parametric Kruskal-Wallis test, followed by Dunn’s multiple comparison post hoc test. *P<0.05, **P<0.01. (f) Frequency distribution of podosome lifetime shown in logarithmic scale. Podosome lifetime was measured by track length analysis of podosomes in 30 min TIRF movies. Note that for depletion of KANK1 or INF2, no long-living podosomes (> 18 min for KANK1 kd; > 24 min for INF2 kd) were observed. For all values, 3 videos of cells per condition cells from each time ≥ 3 donors were analysed. For specific values, see Suppl. Table 2.

**Extended Data Figure 5. Localization of** β **integrin isoforms in primary macrophages.** Micrographs of primary human macrophages stained for endogenous β1 (b,e), β2 (h,k), or β3 (n,q) integrin, with F-actin staining using Alexa568-phalloidin (a,d,g,j,m,p), with merges (c,f,i,l,o,r), analysed by both confocal (a-c,g-i,m-o) and TIRF (d-f,j-l,p-r) microscopy. White boxes in overviews indicate regions of detail images shown in lower right corners. Bars: 10 µm. Note prominent localization of β2 integrin at the podosome ring structure.

**Extended Data Figure 6. AlphaFold3 rendering of electrostatic and hydrophobic interactions between clathrin heavy chain and the LIDL box of various proteins.** (a,b) AF3 model of the CLH1 N-terminus (aa 47-103) in contact with motifs in the drebrin C-terminus, including (a) the canonical clathrin box (aa528-537) or (b) the LIDL box (aa 474-480), with electrostatic surface rendering (positive charge: blue, negative charge: red), with potentially interacting amino acid residues indicated in (c) (see also Fig. 6b). (d-i) AF3 model of the CLH1 N-terminus (aa1-360) in contact with the AAK1 C-terminus (aa 823-961) (ipTM score of 0.68, and pTM score of 0.75). Interacting amino acid residues of CLH1 and AAK1 are colored in magenta and purple respectively, with the LIDL motif in blue letters. Dashed lines represents pseudobonds of interacting atoms with Van der Waals overlap ≥ - 0.1 [. Dashed box in (d) indicates detail region shown enlarged, rotated by 90°, in (e), with (f) further hydrophobicity surface rendering of CHL1 (yellow: hydrophobic; cyan: hydrophilic), or (g) of the AAK1 LIDL box, rotated by 180°, or (h) showing electrostatic surface rendering (positive charge: blue, negative charge: red). (i) Potentially interacting amino acid residues. (j-o) AF3 model of the CLH1 N-terminus (aa1-360) in contact with GSTE1 (aa 1-793) (ipTM score of 0.73 and pTM score of 0.42). Interacting amino acid residues of CLH1 and GSTE1 are colored in magenta and purple, respectively, with the LVDI motif in blue letters Dashed lines represents pseudobonds of interacting atoms with Van der Waals overlap ≥ - 0.1 [. Dashed box indicates detail region shown enlarged, rotated by 90°, in (k), (l) with further hydrophobicity surface rendering of CHL1 (yellow: hydrophobic; cyan: hydrophilic), or (m) of the GTSE1 LVDI box, rotated by 180°, or (n) showing electrostatic surface rendering (positive charge: blue, negative charge: red). (o) Potentially interacting amino acid residues indicated in.

**Extended Data Figure 7. Microtubule posttranslational modifications following INF2 depletion; microtubule +tip heterogeneity;** (a-d) Confocal micrographs of primary macrophages stained for tyrosinated α-tubulin (a,c, green) or acetylated α-tubulin (b,d, green), in cells treated with control siRNA (a,b) or INF2-specific siRNA (c,d), and stained with Alexa 568 for F-actin (red), to detect podosome cores. F-actin and tubulin stainings shown in small side images, with respective merges enlarged. Note that distribution and intensity of either PTM appear unchanged in cells depleted for INF2. Scale bars: 10 µm for overview images, 2 µm for detail images. (e-k) Confocal micrographs of primary macrophages stained for β-tubulin (e, red), and EB3 (g, magenta), and overexpressing CLIP-170-GFP (f, green), with merge (h). (i-k) Merge of only CLIP-170-GFP (green) and EB3, emphasizing their spatial distribution. White boxes indicate areas of detail images shown in (j,k). Colored arrowheads indicate +tips showing predominantly CLIP-170-GFP, predominantly EB3, or mixed signals. Respective single channels are shown on right. Scale bars: 5 µm for whole-cell images and 2 µm for detail images.

**Extended Data Figure 8. Colocalization of clathrin and EEA1 in a polarized macrophage.** (a) Maximum projections of confocal micrographs of primary human macrophages, stained with respective antibodies for endogenous CLH1 (green), or EEA1 (red), and overexpressing drebrin-GFP (magenta), and stained with Alexa405-phalloidin for F-actin (white). (b) Different confocal planes of the cell in (a) are shown, starting from the top (Z=4) to the ventral side of podosomes (Z=0), with (c) showing colocalizing pixels of CLH1 and EEA1 (cyan). Note colocalization of CLH1 and EEA1 at higher optical planes (Z=4 to Z=2) in podosome-containing regions and at lower optical planes (Z=1) in regions devoid of podosomes, likely reflecting colocalization at the podosome cap or the cell cortex, respectively. Scale bar: 10 µm. See also (Fig. 5r-w).

**Extended Data Figure 9. AlphaFold3 model of drebrin binding to actin nuclei or filaments of increasing length.** Drebrin (color coded blue to red from N- to C-terminus) in the presence of a nucleus of 3 actin monomers (grey), an actin filament of 4 actin monomers, or an actin filament of 8 monomers. Note how the C-terminus of drebrin (yellow to red) is normally unstructured, but it folds when bound to an actin filament. As this region contains the clathrin-binding motifs, binding of drebrin to F-actin could also affect its binding to clathrin heavy chain. Dashed white boxed indicate the canonical clathrin binding box “LLNFDEL” harboured in the C-terminus of drebrin.

## Supplementary Videos

**Suppl. Video 1. EB3-GFP-positive MT +tips contact drebrin-RFP-positive podosome caps** 4D-SoRa time lapse videos of a macrophage overexpressing Drebrin-RFP (red) together with EB3-GFP (green). Images were acquired as 3D stacks over time using the Visitron microscope in super-resolution mode (SoRa module). Videos show on the left the bottom Z-slice (as in fig. 3s), in the center the middle Z-slice (as in fig. 3q) and on the right the top Z-slice (as in fig. 3o). Note how most contacts between EB3 and podosomes occur in the middle-to-top part of podosomes. 100X objective, 5 sec / frame. See (fig. 3 o-t).

**Suppl. Video 2. EB3-positive MT +tips contact drebrin-positive podosome caps** SoRa time lapse videos of a macrophage overexpressing Drebrin-RFP (red) together with EB3-GFP (green). Images were acquired using the Visitron microscope in super-resolution mode (SoRa module), focusing only on the middle Z-plane of podosomes. 100X objective, 1 sec / frame. See (fig. 3u).

**Suppl. Video 3. EB3-positive MT +tips contact drebrin-positive podosome caps.** ROI of suppl. Video 2. Note how the EB3-decorated +tip has prolonged, likely multiple, contact with the same podosome. See (fig. 3w).

**Suppl. Video 4. EB3-positive MT +tips contact drebrin-positive podosome caps.** ROI of suppl. Video 2. Note how the EB3 decorated microtubule +tip “slides” between podosomes. See (fig. 3x).

**Suppl. Video 5. Clathrin light chain-positive vesicles contact the drebrin-positive podosome cap.** SoRa time lapse videos of a macrophage overexpressing Drebrin-RFP (red) together with clathrin light chain-GFP (green). Images were acquired using the Visitron microscope in super-resolution mode (SoRa module), focusing only on the middle-top Z-plane of podosomes. 100X objective, 5 sec / frame. See (fig. 5x)

**Suppl. Video 6. Clathrin light chain-positive vesicles contact the drebrin-positive podosome cap.** ROI of suppl. Video 5; 100X objective, 5 sec / frame. See (Fig. 5x1).

**Suppl. Video 7. Clathrin light chain-positive vesicles contact the drebrin-positive podosome cap.** ROI of suppl. Video 5; 100X objective, 5 sec / frame. See (Fig. 5x2).

**Suppl. Video 8. Drebrin conformation is affected by actin filament length.** 360° animation of AlphaFold 3 modelling of drebrin full length, color-coded rainbow from the N-terminus (blue) to C-terminus (red), together with 3 (left), 4 (center) or 8 (right) G-actin monomers structured as F-actin, colered in light gray. The white letters indicate the position of the canonical clathrin binding box “LLNFDEL”. Note how the 3D conformation of the C-terminus of drebrin is affected by the length of the actin filament, determining the availability of the canonical clathrin binding box to bind clathrin. See (Ext. Data Fig.9).

