## Supplementary material for "Drebrin forms a cortical hub that connects actin, microtubules, and clathrin for endocytic transport of β2 integrin": Suppl. Tables 1-3

**Suppl. Table 1.** Used siRNAs, cloning primers and constructs

| Used siRNA | | |
|---|---|---|
| **Target** | **Sequence** | **Reference** |
| ctrl | 5'- AGGUAGUGUAACCGCCUUGUU -3' | 10 |
| IQGAP1 | 5'- GAACGUGGCUUAUGAGUAC -3' | Dharmacon. Ref#: SO-2734064G |
| KANK1 | human KANK1 #1: 5´-CAGAGAAGGACATGCGGAT-3´<br>human KANK1 #2: 5´-GAAGTCAGCGTCTGCGAAA-3´ | 38 |
| ELKS | 5´-GTGGGAAAACCCTTTCAAT-3´ | 37 |
| PAK4 | 5′-GGTGAACATGTATGAGTGT-dtdt-3′ | Walls et al., J. Cell Sci, 2010 |
| SV | 5'- CAGCCAUAAGGAAUCUAAAUAUGCU -3’ | 22 |
| INF2 | 5'- CCAUGAAGGCUUUCCGGGA -3' | 26 |
| LSP1 | 5′- UGGAGACAUGAGCAAGAAA -3′ | 24 |
| DBN1 | 5'- GGAAACAGCAGACUUUAGA -3' | 52 |
| EB3 | 5´-CAGCAAACUUCGUGACAUC-dtdt-3´ | 56 |
| EB1 | 5'- UGACAAAGAUCGAACAGUU -3' | 37 |

| Used primer for cloning | | |
|---|---|---|
| **Construct** | **Template** | **Sequence [5’-3’]** |
| ΔADFH-GFP | DBN1-GFP | rev:[PHO]CATGGCGGCCGCTAGCGGATCTGACGGTTCACTAAAC<br>fw: [PHO]GGGCTGGCGCGACTCTCC |
| ΔADFH-ΔPP-GFP | ΔADFH-GFP | rev: [PHO]CTGCGAGGAGGTGACCTCATCC<br>fw: [PHO]GCAGAGGACTTGATGTTCATGGAGTCTGC |
| CC-Hel-GFP | ΔADFH-GFP | rev: [PHO]CTGCGAGGAGGTGACCTCATCC<br>fw: [PHO]CCGGTCGCCACCATGGTGAG |
| PP-C-term-GFP | DBN1-GFP | rev:[PHO]CATGGCGGCCGCTAGCGGATCTGACGGTTCACTAAAC<br>fw: [PHO]CCTCCACCACTGCCACCGCC |
| C-term-GFP | DBN1-GFP | rev: [PHO]CATGGCGGCCGCTAGCGGATCTGACGGTTCACTAAAC<br>fw: [PHO]GCAGAGGACTTGATGTTCATGGAGTCTGC |

| Used constructs | |
|---|---|
| **Construct** | **Reference** |
| EB3-GFP | Marina Mikhaylova |
| EB3-mRFP | Niels Galjart |
| EB1-GFP | Yuko Mimori-Kiyosue |
| LifeAct-RFP | Michael Sixt |
| CLIP170-GFP | Niels Galjart |
| DBN1-GFP | in house |
| mCherry ctrl | in house |
| DBN1 constructs | in house |

**Suppl. Table 2.** Values from measurements

| Figure 3 a-f: Quantification of contact parameters with *ContactAnalyzer* | | | | |
|---|---|---|---|---|
| | **siRNA target** | **Quantification (Mean. ±SD; ± S.E.M.)** | | |
| a) Podosome lifetime | ctrl | 3.8 | 1.187 | 0.1662 |
| | IQGAP1 | 3.946 | 1.229 | 0.4095 |
| | KANK1 | 2.291 | 0.8622 | 0.3048 |
| | ELKS | 4.167 | 1.513 | 0.4198 |
| | PAK4 | 3.648 | 1.268 | 0.3517 |
| | SV | 3.711 | 0.7639 | 0.2119 |
| | INF2 | 2.834 | 1.078 | 0.3595 |
| | LSP1 | 4.647 | 0.8422 | 0.2978 |
| | DBN1 | 2.959 | 1.13 | 0.3767 |
| b) Cumulative podosome number | ctrl | 1508 | 600.1 | 83.22 |
| | IQGAP1 | 1525 | 499.3 | 166.4 |
| | KANK1 | 1540 | 1215 | 429.4 |
| | ELKS | 1470 | 553.8 | 153.6 |
| | PAK4 | 1272 | 519.3 | 144 |
| | SV | 1202 | 299.6 | 83.08 |
| | INF2 | 867,3 | 679.3 | 226.4 |
| | LSP1 | 1512 | 556.3 | 185.4 |
| | DBN1 | 1531 | 661.2 | 220.4 |
| c) Raw contacts | ctrl | 5530 | 4301 | 596.5 |
| | IQGAP1 | 10264 | 7166 | 2389 |
| | KANK1 | 6307 | 8288 | 2930 |
| | ELKS | 6568 | 5085 | 1410 |
| | PAK4 | 3269 | 3146 | 872.5 |
| | SV | 3830 | 2113 | 586.1 |
| | INF2 | 8521 | 7362 | 2454 |
| | LSP1 | 4028 | 2373 | 791 |
| | DBN1 | 883.4 | 643.4 | 227.5 |
| d) Consecutive contacts | ctrl | 4507 | 3492 | 484.2 |
| | IQGAP1 | 8357 | 5591 | 1864 |

| | | | | |
|---|---|---|---|---|
| | KANK1 | 5587 | 7337 | 2594 |
| | ELKS | 5707 | 4377 | 1214 |
| | PAK4 | 2523 | 2321 | 643.8 |
| | SV | 3352 | 1825 | 506.1 |
| | INF2 | 6100 | 5564 | 1855 |
| | LSP1 | 3588 | 1943 | 647.8 |
| | DBN1 | 760.4 | 551.4 | 195 |
| e) Contacts/podosome | ctrl | 3.378 | 2.915 | 0.4043 |
| | IQGAP1 | 5.7 | 3.462 | 1.154 |
| | KANK1 | 2.814 | 2.541 | 0.8983 |
| | ELKS | 3.89 | 2.372 | 0.658 |
| | PAK4 | 1.997 | 1.395 | 0.3868 |
| | SV | 2.838 | 1.46 | 0.4049 |
| | INF2 | 6.16 | 3.152 | 1.051 |
| | LSP1 | 2.752 | 1.897 | 0.6323 |
| | DBN1 | 0.4843 | 0.2741 | 0.0969 |
| f) Contact duration (seconds) | ctrl | 2.472 | 0.3496 | 0.04848 |
| | IQGAP1 | 2.458 | 0.3541 | 0.118 |
| | KANK1 | 2.256 | 0.1067 | 0.03773 |
| | ELKS | 2.294 | 0.1753 | 0.04861 |
| | PAK4 | 2.588 | 0.3787 | 0.105 |
| | SV | 2.271 | 0.1501 | 0.04164 |
| | INF2 | 2.815 | 0.341 | 0.1137 |
| | LSP1 | 2.19 | 0.142 | 0.04732 |
| | DBN1 | 2.326 | 0.1664 | 0.05883 |

| **Figure 3m: Quantification of EB3/DBN1-GFP PLA signals** | | |
|---|---|---|
| **Staining** | **Cell number** | **Signals/cell (Mean; ±SD; ±S.E.M.)** |
| ctrl | 30 | 10.47 ± 6.637 |
| EB3/DBN1-GFP | 31 | 364.7 ± 146.8 |

| Figure 3n: Quantification of relative signal height and distance to podosome center | | |
|---|---|---|
| **Number of signals** | **Median. ±SD** | |
| 1.506 | Distance to podosome centroid [µm] | 0.49 ± 0.17 |
| | relative signal height [µm] | 0.86 ± 0.24 |

| Figure 4b: Streptavidin norm. fl. int. [%] | | | |
|---|---|---|---|
| **siRNA** | **Time [min]** | **Mean value/%** | **±SD** |
| ctrl | surf | 10515 / 100 | ± 10515 |
| | 10 | 11327 / 48.004 | ± 11327 |
| | 30 | 5683 / 29.72 | ± 5683 |
| | 60 | 7572 / 26.77 | ± 7572 |
| | 120 | 5318 / 18.39 | ± 5318 |
| DBN1 | surf | 43016 / 100.39 | ± 13868 |
| | 10 | 12157 / 28.37 | ± 6676 |
| | 30 | 6273 / 14.64 | ± 4540 |
| | 60 | 3320 / 7.75 | ± 2203 |
| | 120 | 3606 / 8.42 | ± 2236 |

| Figure 4d-f: Quantification of the surface biotinylation assay | | | |
|---|---|---|---|
| **Target** | **Time [min]** | **ctrl siRNA (Mean; ±SD)** | **DBN1 siRNA (Mean; ±SD)** |
| ITGB1 (n=2) | 0 | 100 ± 1.21 | 100 ± 0.38 |
| | 30 | 29.5 ± 4.11 | 37.09 ± 8.26 |
| | 60 | 14.56 ± 6.21 | 7.81 ± 1.42 |
| | 120 | 16.90 ± 0.14 | 11.91 ± 7.64 |
| ITGB2 (n=2) | 0 | 100 ± 8.74 | 100 ± 15.65 |
| | 30 | 75.98 ± 2.33 | 74.39 ± 4.86 |
| | 60 | 59.30 ± 2.01 | 47.87 ± 4.52 |
| | 120 | 82.81 ± 10.37 | 51.26 ± 8.35 |
| ITGB3 (n=2) | 0 | 100 ± 16.81 | 100 ± 15.17 |
| | 30 | 49.79 ± 3.02 | 60.04 ± 6.64 |
| | 60 | 29.79 ± 1.68 | 34.28 ± 10.52 |

| | | | |
|---|---|---|---|
| | 120 | 41.05 ± 9.43 | 37.66 ± 10.76 |

| Figure 4g: Quantification of endogenous target protein expression after DBN1 depletion | | | |
|---|---|---|---|
| **siRNA target** | **Sample no.** | **Expression (mean)** | **±SD; ±S.E.M.** |
| ctrl (luc) | | set to 100 ea. | ± 0; ± 0 |
| ITGB1 | 2 | 96.50 | ± 9.19; ± 6.5 |
| ITGB2 | 5 | 98.00 | ± 6.29; ± 2.81 |
| ITGB3 | 3 | 89.00 | ± 18.03; ± 10.41 |

| Figure 5p: Quantification of CLH1/DBN1 PLA signals | | |
|---|---|---|
| **Staining** | **Cell number** | **Signals/cell (Mean; ±SD)** |
| ctrl | 158 | 3.918 ± 4.864 |
| CLH1/DBN1 | 57 | 493.2 ± 231.7 |

| Figure 5: Quantification of relative signal height and distance to podosome center | | |
|---|---|---|
| **Number of signals** | **Median. ±SD** | |
| 1.748 | distance to podosome centroid [µm] | 0.48 ± 0.13 |
| | relative signal height [µm] | 1.00 ± 0.26 |

| Figure 6f: Quantification of GFP IP | | |
|---|---|---|
| **Construct** | **Values** | **Mean. ± SD** |
| EGFP ctrl | n=3 | 38.67 ± 24 |
| DBN1-PP + C-term | n=3 | 100 ± 0.0 |
| DBN1-PP + C-term #1 | n=3 | 89 ± 20 |
| DBN1-PP + C-term #1+2 | n=3 | 35 ± 6.9 |

| Ext. Data Figure 2d-i: Quantification of contact parameters with *ContactAnalyzer* | | | | |
|---|---|---|---|---|
| **Parameter** | **Overexpression** | **Quantification (Mean. ±SD; ± S.E.M.)** | | |
| podosomes (number per cell) | CLIP170 | 846.3 | 426.1 | 113.9 |
| | EB1 | 1006 | 576.9 | 192.3 |
| | EB3 | 942.3 | 361.1 | 104.2 |
| MT+ Tips | CLIP170 | 87302 | 63771 | 17044 |

| | | | | |
|---|---|---|---|---|
| (number per cell) | EB1 | 15728 | 17923 | 5974 |
| | EB3 | 57984 | 43796 | 12643 |
| raw contacts | CLIP170 | 46692 | 27511 | 7353 |
| | EB1 | 13008 | 13786 | 4595 |
| | EB3 | 26337 | 18256 | 5270 |
| consecutive contacts | CLIP170 | 26962 | 17844 | 4769 |
| | EB1 | 8927 | 8134 | 2711 |
| | EB3 | 10441 | 7181 | 2073 |
| contacts per podosome | CLIP170 | 32.36 | 17.84 | 4.768 |
| | EB1 | 9.083 | 7.906 | 2.635 |
| | EB3 | 11.05 | 7.082 | 2.045 |
| contact duration (seconds) | CLIP170 | 3.722 | 0.9518 | 0.2544 |
| | EB1 | 2.603 | 0.4415 | 0.1472 |
| | EB3 | 5.159 | 1.307 | 0.3774 |

| **Ext. Data Figure 4b-f: Quantification of cell parameters with *ContactAnalyzer*** | | | | |
|---|---|---|---|---|
| | **siRNA target** | **Quantification (Mean. ±SD; ± S.E.M.)** | | |
| b) cell area | ctrl | 419 | 165 | 22 |
| | IQGAP1 | 506 | 184 | 61 |
| | KANK1 | 375 | 116 | 41 |
| | ELKS | 339 | 146 | 39 |
| | PAK4 | 408 | 111 | 28 |
| | SV | 305 | 96 | 27 |
| | INF2 | 210 | 88 | 31 |
| | LSP1 | 381 | 164 | 39 |
| | DBN1 | 493 | 104 | 35 |
| c) Podosomes (number/cell) | ctrl | 317 | 131 | 17 |
| | IQGAP1 | 340 | 128 | 43 |
| | KANK1 | 213 | 127 | 45 |
| | ELKS | 297 | 120 | 32 |
| | PAK4 | 296 | 101 | 25 |
| | SV | 223 | 52 | 15 |
| | INF2 | 168 | 86 | 30 |

| | | | | |
|---|---|---|---|---|
| | LSP1 | 340 | 163 | 38 |
| | DBN1 | 397 | 102 | 34 |
| d) Podosome density | ctrl | 0.77 | 0.18 | 0.02 |
| | IQGAP1 | 0.68 | 0.11 | 0.04 |
| | KANK1 | 0.54 | 0.18 | 0.06 |
| | ELKS | 0.89 | 0.10 | 0.03 |
| | PAK4 | 0.72 | 0.14 | 0.03 |
| | SV | 0.77 | 0.19 | 0.05 |
| | INF2 | 0.79 | 0.16 | 0.06 |
| | LSP1 | 0.90 | 0.15 | 0.04 |
| | DBN1 | 0.82 | 0.21 | 0.07 |
| e) MT+ Tips (number/cell) | ctrl | 23021 | 14139 | 1961 |
| | IQGAP1 | 53503 | 26756 | 8919 |
| | KANK1 | 41982 | 35112 | 12414 |
| | ELKS | 25625 | 22666 | 6286 |
| | PAK4 | 17607 | 12496 | 3466 |
| | SV | 15431 | 6871 | 1906 |
| | INF2 | 34853 | 13935 | 4645 |
| | LSP1 | 15073 | 9481 | 3160 |
| | DBN1 | 15037 | 17353 | 6135 |
| f) Frequency (min) | ctrl | 8.5 | 51 | 14 |
| | IQGAP1 | 8.6 | 53 | 15 |
| | KANK1 | 6.48 | 37 | 10 |
| | ELKS | 8.66 | 50 | 14 |
| | PAK4 | 8.21 | 40 | 11 |
| | SV | 7.67 | 45 | 12 |
| | INF2 | 7.28 | 27 | 7 |
| | LSP1 | 9.39 | 58 | 16 |
| | DBN1 | 7.66 | 47 | 13 |

**Suppl. Table 3** Identified proteins containing the LIDL motif

| uniprot id | gene name | full name | motif start | motif end | motif [L,M,I,V] [I,F,V,M] D [L,F,I,M] | mask |
|---|---|---|---|---|---|---|
| Q2M2I8 | AAK1 | AP2-associated protein kinase 1 | 958 | 961 | L L V D Q **L I D L** - - - - - | LIDL |
| Q16643 | DBN1 | Drebrin | 475 | 478 | P A A T S **L I D L** W P G N G | LIDL |
| Q9NZ52 | GGA3 | ADP-ribosylation factor-binding protein GGA3 | 326 | 329 | S N Q G T **L I D L** A E L D T | LIDL |
| Q9NYZ3 | GTSE1 | G2 and S phase-expressed protein 1 | 694 | 697 | P V V G Q **L I D L** S S P L I | LIDL |
| Q9NYZ3 | GTSE1 | G2 and S phase-expressed protein 1 | 645 | 648 | A A S Q P **L I D L** P L I D F | LIDL |
| Q96D71 | REPS1 | RalBP1-associated Eps domain-containing protein 1 | 371 | 374 | S L M P K **L I D L** E D S A D | LIDL |
| P11274 | BCR | Breakpoint cluster region protein | 260 | 263 | F L K D N **L I D A** N G G S R | LID- |
| Q8N684 | CPSF7 | Cleavage and polyadenylation specificity factor subunit 7 | 7 | 10 | S E G V D **L I D I** Y A D E E | LID- |
| Q9NYZ3 | GTSE1 | G2 and S phase-expressed protein 1 | 650 | 653 | L I D L P **L I D F** C D T P E | LID- |
| Q01968 | OCRL | Inositol polyphosphate 5-phosphatase OCRL | 73 | 76 | A E E T L **L I D I** A S N S G | LID- |
| O15027 | SEC16A | Protein transport protein Sec16A | 1579 | 1582 | P N E A N **L I D F** T N E A V | LID- |
| Q8N3V7 | SYNPO | Synaptopodin | 437 | 440 | M S S S L **L I D I** Q P N T L | LID- |
| Q07912 | TNK2 | Activated CDC42 kinase 1 | 570 | 573 | G A E V T **L I D F** G E E P V | LID- |
| O60784 | TOM1 | Target of Myb1 membrane trafficking protein | 322 | 325 | E P A A D **L I D M** G P D P A | LID- |
| Q8TES7 | FBF1 | Fas-binding factor 1 | 919 | 922 | Q R E G T **L I S L** A K Q A E | LI-L |
| Q9ULH0 | KIDINS220 | Kinase D-interacting substrate of 220 kDa | 1614 | 1617 | L E K A N **L I E L** E D D S H | LI-L |
| O43426 | SYNJ1 | Synaptojanin-1 | 1355 | 1358 | P D P K R **L I Q L** P S A T Q | LI-L |
| Q69YN4 | VIRMA | Protein virilizer homolog | 1552 | 1555 | L V K V D **L I E L** S E K C C | LI-L |
| Q10567 | AP1B1 | AP-1 complex subunit beta-1 | 702 | 705 | S G L S D **L F D L** T S G V G | L-DL |
| Q676U5 | ATG16L1 | Autophagy-related protein 16-1 | 152 | 155 | D L R T K **L C D L** E R A N Q | L-DL |
| Q14677 | CLINT1 | Clathrin interactor 1 | 326 | 329 | K S S G D **L V D L** F D G T S | L-DL |
| Q14677 | CLINT1 | Clathrin interactor 1 | 423 | 426 | S N S S D **L F D L** M G S S Q | L-DL |
| P98082 | DAB2 | Disabled homolog 2 | 236 | 239 | S K D I L **L V D L** N S E I D | L-DL |
| Q09472 | EP300 | Histone acetyltransferase p300 | 36 | 39 | T D F G S **L F D L** E H D L P | L-DL |
| Q9Y6I3 | EPN1 | Epsin-1 | 257 | 260 | K E E S S **L M D L** A D V F T | L-DL |
| Q9Y6I3 | EPN1 | Epsin-1 | 481 | 484 | G P N A A **L V D L** D S L V S | L-DL |
| O00165 | HAX1 | HCLS1-associated protein X-1 | 3 | 6 | - - - M S **L F D L** F R G F F | L-DL |
| Q9HAU0 | PLEKHA5 | Pleckstrin homology domain-containing family A member 5 | 383 | 386 | I V N V S **L A D L** R G G N R | L-DL |
| P49023 | PXN | Paxillin | 8 | 11 | D L D A L **L A D L** E S T T S | L-DL |
| Q9HBD1 | RC3H2 | Roquin-2 | 866 | 869 | N S N A V **L M D L** D S G D V | L-DL |
| O60641 | SNAP91 | Clathrin coat assembly protein AP180 | 898 | 901 | P A K D P **L A D L** N I K D F | L-DL |
| O95782 | AP2A1 | AP-2 complex subunit alpha-1 | 682 | 685 | G A G N L **L V D V** F D G P A | L-D- |
| O94973 | AP2A2 | AP-2 complex subunit alpha-2 | 684 | 687 | G G S G L **L V D V** F S D S A | L-D- |
| O75061 | DNAJC6 | Putative tyrosine-protein phosphatase auxilin | 22 | 25 | S Y G G G **L F D M** V K G G A | L-D- |
| O60469 | DSCAM | Cell adhesion molecule DSCAM | 1718 | 1721 | Q A T G P **L V D V** S D A R P | L-D- |
| Q9NYZ3 | GTSE1 | G2 and S phase-expressed protein 1 | 571 | 574 | K T D S R **L V D V** S P D R G | L-D- |
| Q9NYZ3 | GTSE1 | G2 and S phase-expressed protein 1 | 626 | 629 | P S E A L **L V D I** K L E P L | L-D- |
| P04150 | NR3C1 | Glucocorticoid receptor | 62 | 65 | K Q R R L **L V D F** P K G S V | L-D- |
| P29353 | SHC1 | SHC-transforming protein 1 | 458 | 461 | S A P R D **L F D M** K P F E D | L-D- |
| Q15398 | DLGAP5 | Disks large-associated protein 5 | 60 | 63 | L E G R I **L V E L** D E T S Q | L--L |
| O95208 | EPN2 | Epsin-2 | 519 | 522 | G P N A A **L V N L** D S L V T | L--L |
| O14617 | AP3D1 | AP-3 complex subunit delta-1 | 706 | 709 | H I P V V **Q I D L** S V P L K | -IDL |
| Q14677 | CLINT1 | Clathrin interactor 1 | 284 | 287 | A N P S K **T I D L** G A A A H | -IDL |
| Q8N684 | CPSF7 | Cleavage and polyadenylation specificity factor subunit 7 | 27 | 30 | F N N T D **Q I D L** Y D D V L | -IDL |
| Q8TC44 | POC1B | POC1 centriolar protein homolog B | 351 | 354 | N P K L E **V I D L** Q I S T P | -IDL |
| Q14498 | RBM39 | RNA-binding protein 39 | 396 | 399 | A E F S F **V I D L** Q T R L S | -IDL |
| P12270 | TPR | Nucleoprotein TPR | 1917 | 1920 | T S Q S L **Q I D L** G P L Q S | -IDL |
| Q96SN8 | CDK5RAP2 | CDK5 regulatory subunit-associated protein 2 | 1 | 4 | - - - - - **M M D L** V L E E D | --DL |
| Q969H0 | FBXW7 | F-box/WD repeat-containing protein 7 | 149 | 152 | T N S S S **I V D L** P V H Q L | --DL |
| O00165 | HAX1 | HCLS1-associated protein X-1 | 267 | 270 | D D A F S **I L D L** F L G R W | --DL |
| O60303 | KATNIP | Katanin-interacting protein | 1142 | 1145 | F Y S D E **M F D L** D V G S L | --DL |
| Q5T7N2 | L1TD1 | LINE-1 type transposase domain-containing protein 1 | 40 | 43 | K D I A P **V L D L** K C K D V | --DL |
| Q9HAU0 | PLEKHA5 | Pleckstrin homology domain-containing family A member 5 | 875 | 878 | R K T K K **M M D L** R T E R P | --DL |
| Q8NA72 | POC5 | Centrosomal protein POC5 | 162 | 165 | Q K M E N **V L D L** W S S G L | --DL |
| P78317 | RNF4 | E3 ubiquitin-protein ligase RNF4 | 46 | 49 | T A G D E **I V D L** T C E S L | --DL |
| P78317 | RNF4 | E3 ubiquitin-protein ligase RNF4 | 58 | 61 | S L E P V **V V D L** T H N D S | --DL |
| Q6ZNL6 | FGD5 | FYVE, RhoGEF and PH domain-containing protein 5 | 612 | 615 | S T P S S **M V D I** P P P F D | --D- |
