## Supplementary material for "Drebrin forms a cortical hub that connects actin, microtubules, and clathrin for endocytic transport of β2 integrin": Ext Data Fig 1-9

**Extended Data Fig. 1**

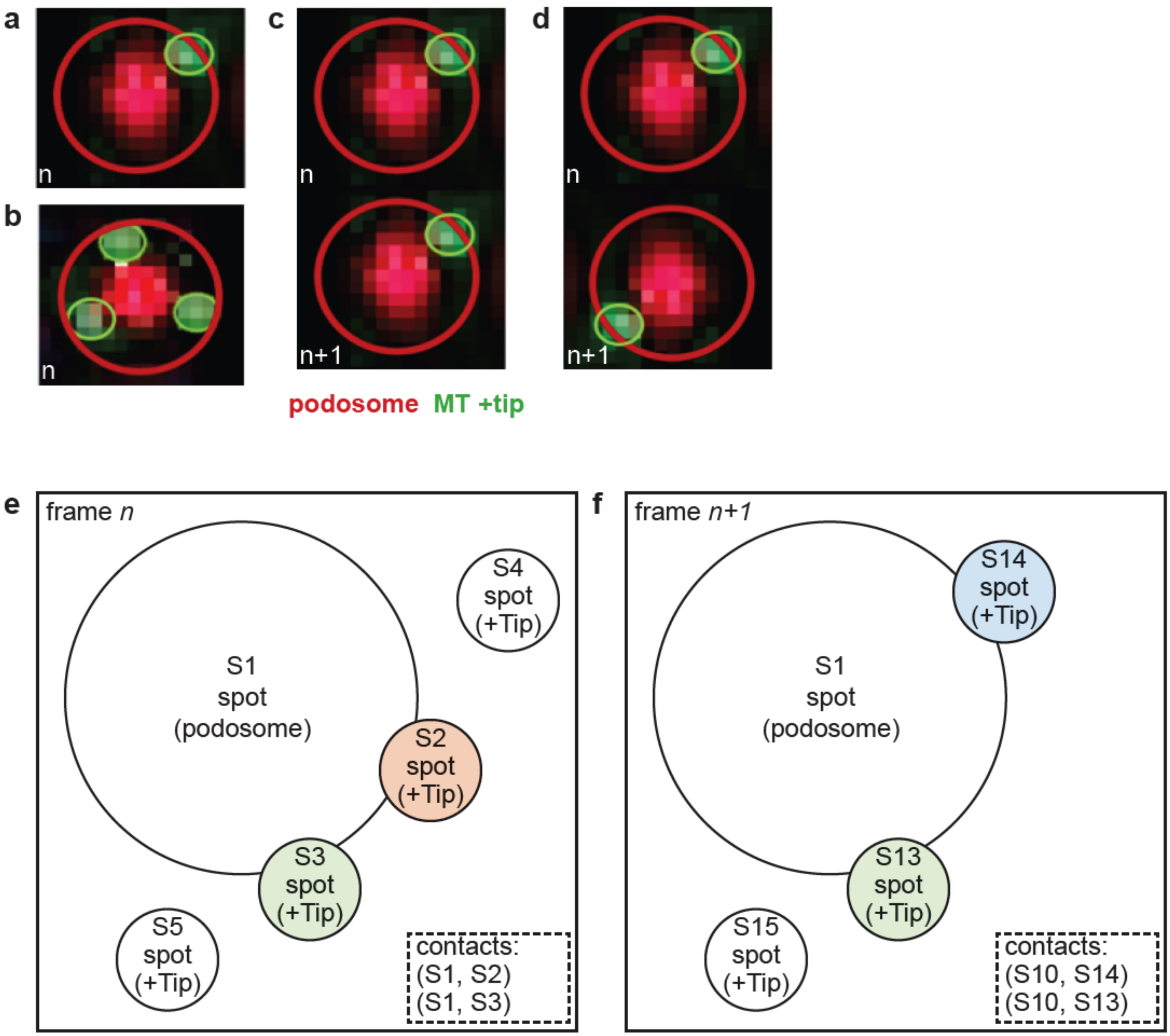

**Extended Data Fig. 2**

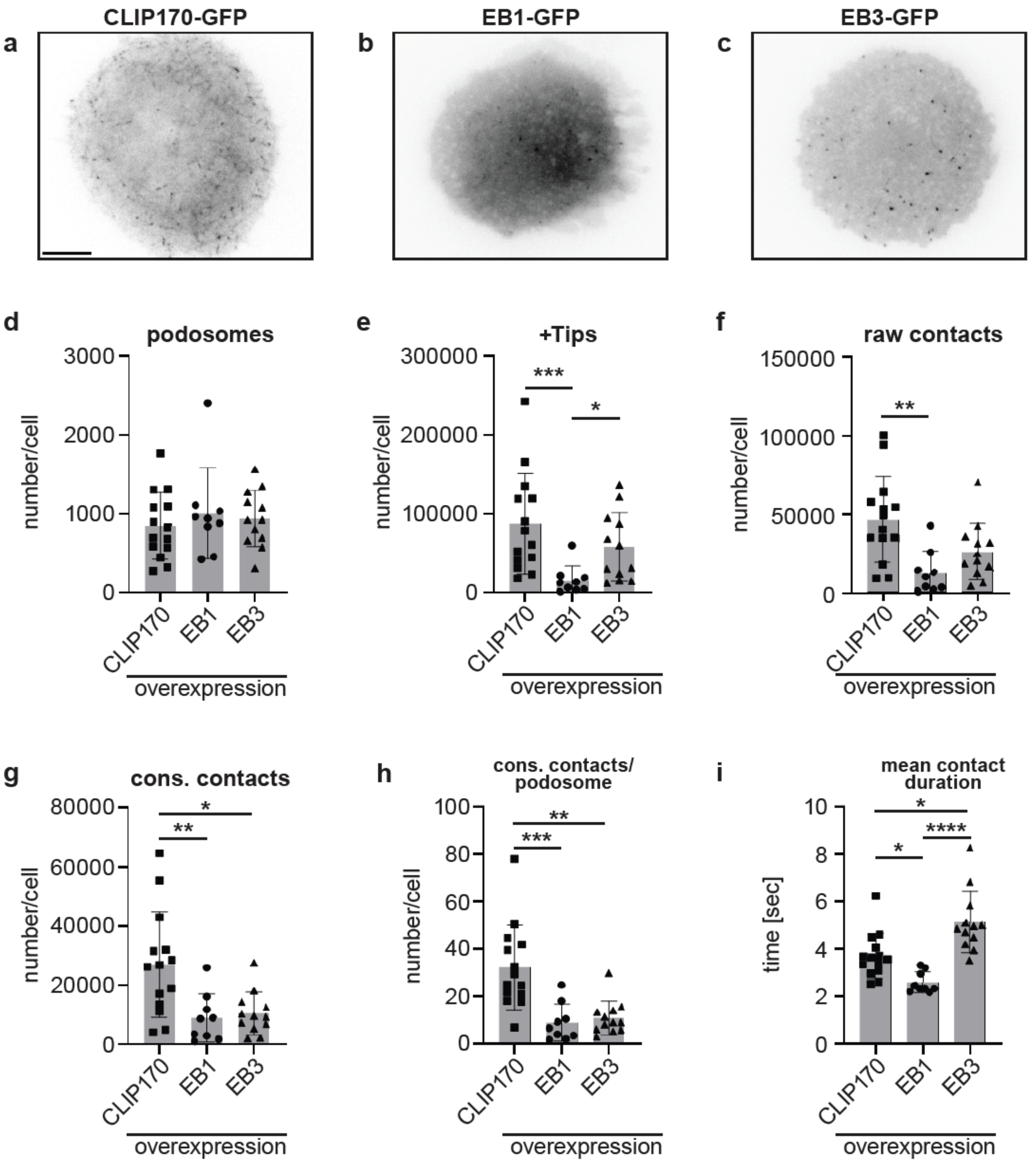

### Extended Data Fig. 3

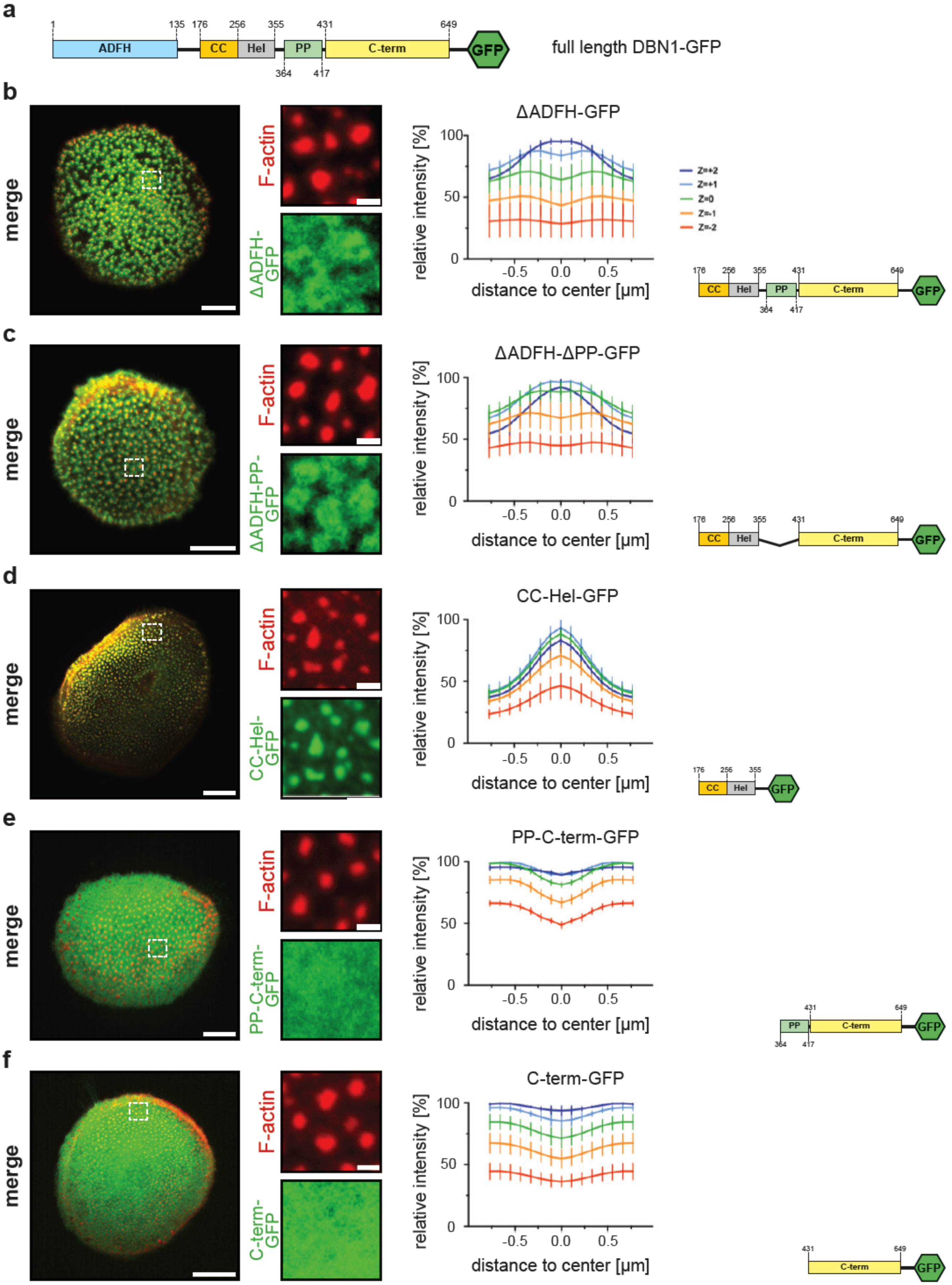

**Extended Data Fig. 4**

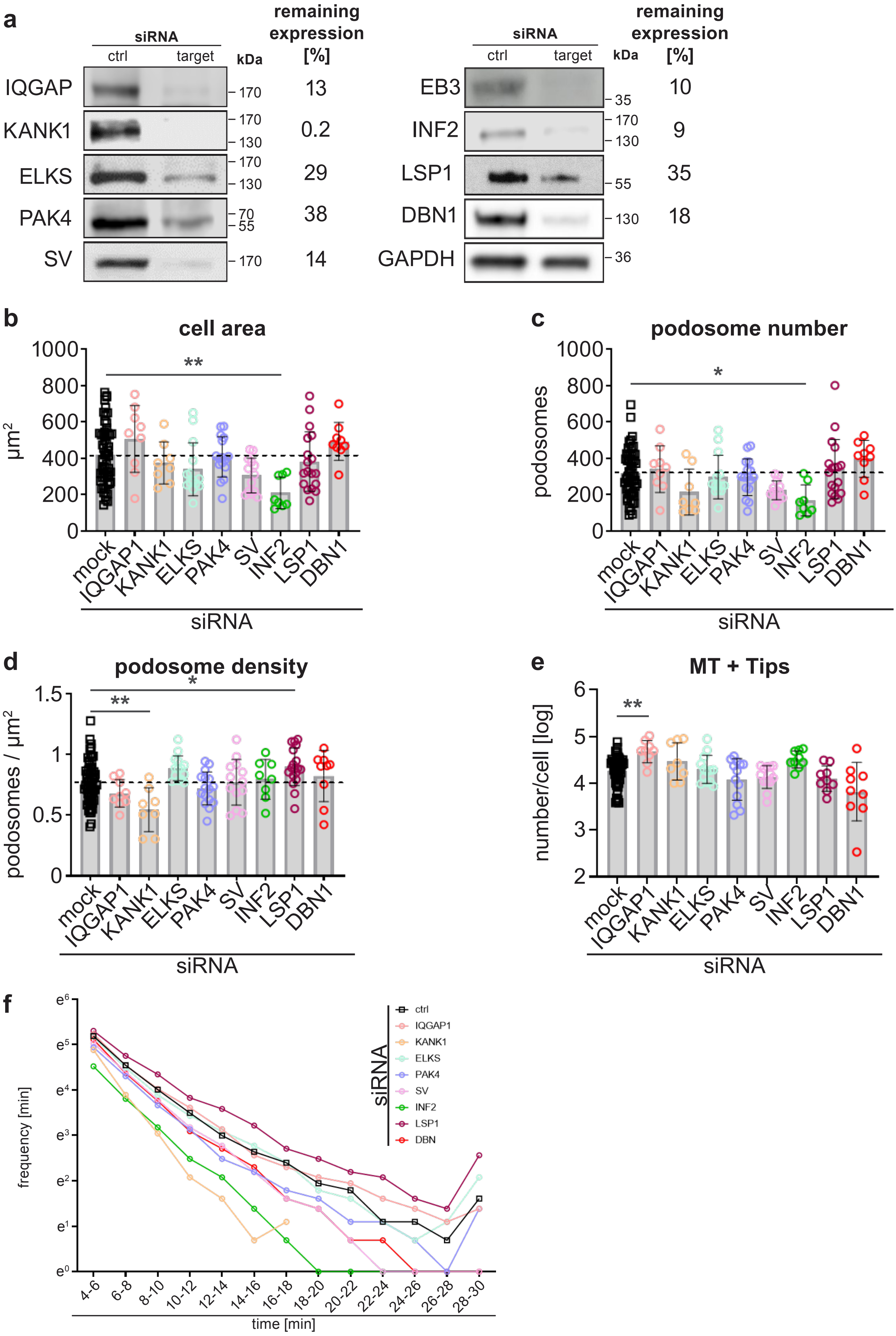

### Extended Data Fig. 5

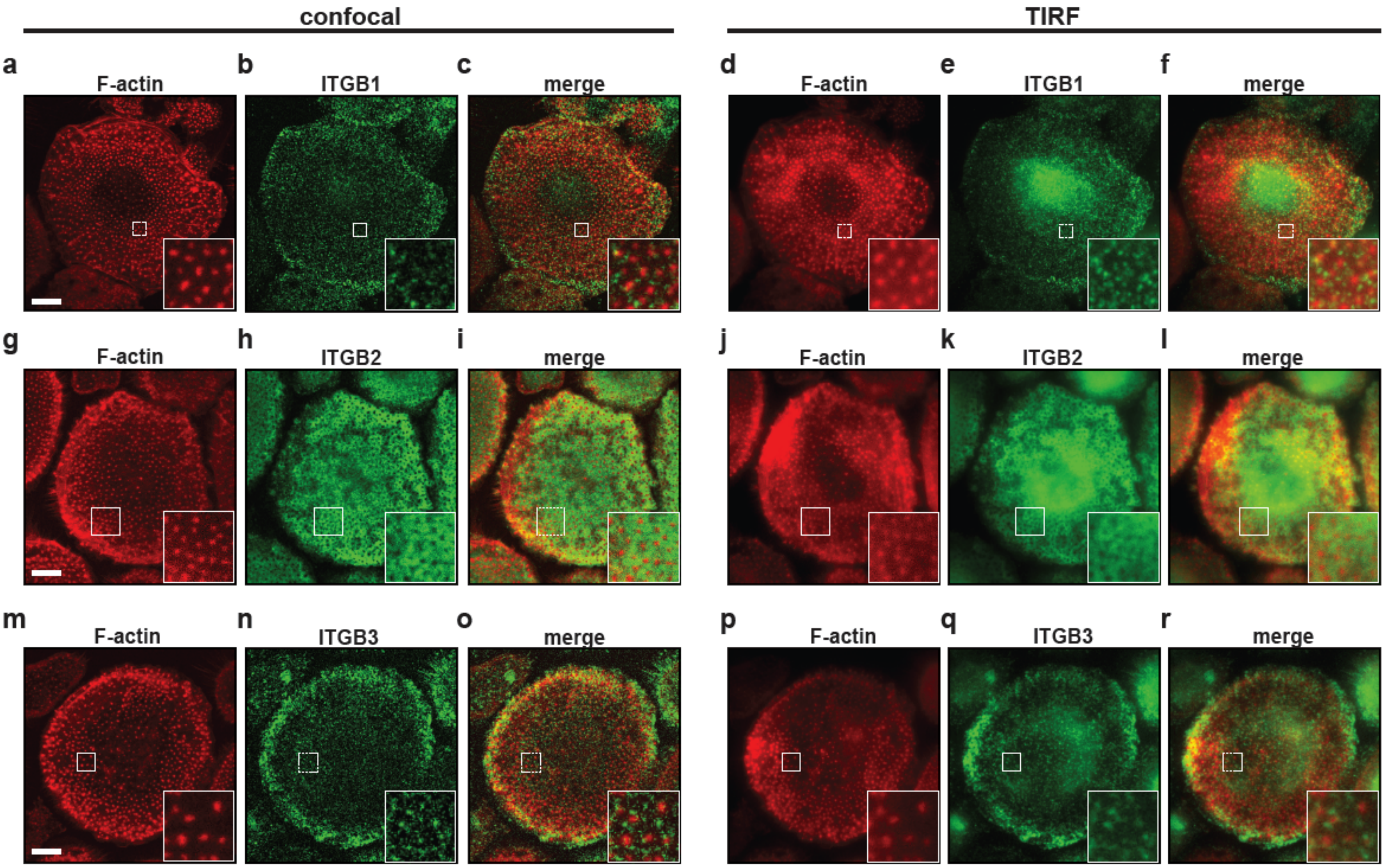

**Extended Data Fig. X 6**

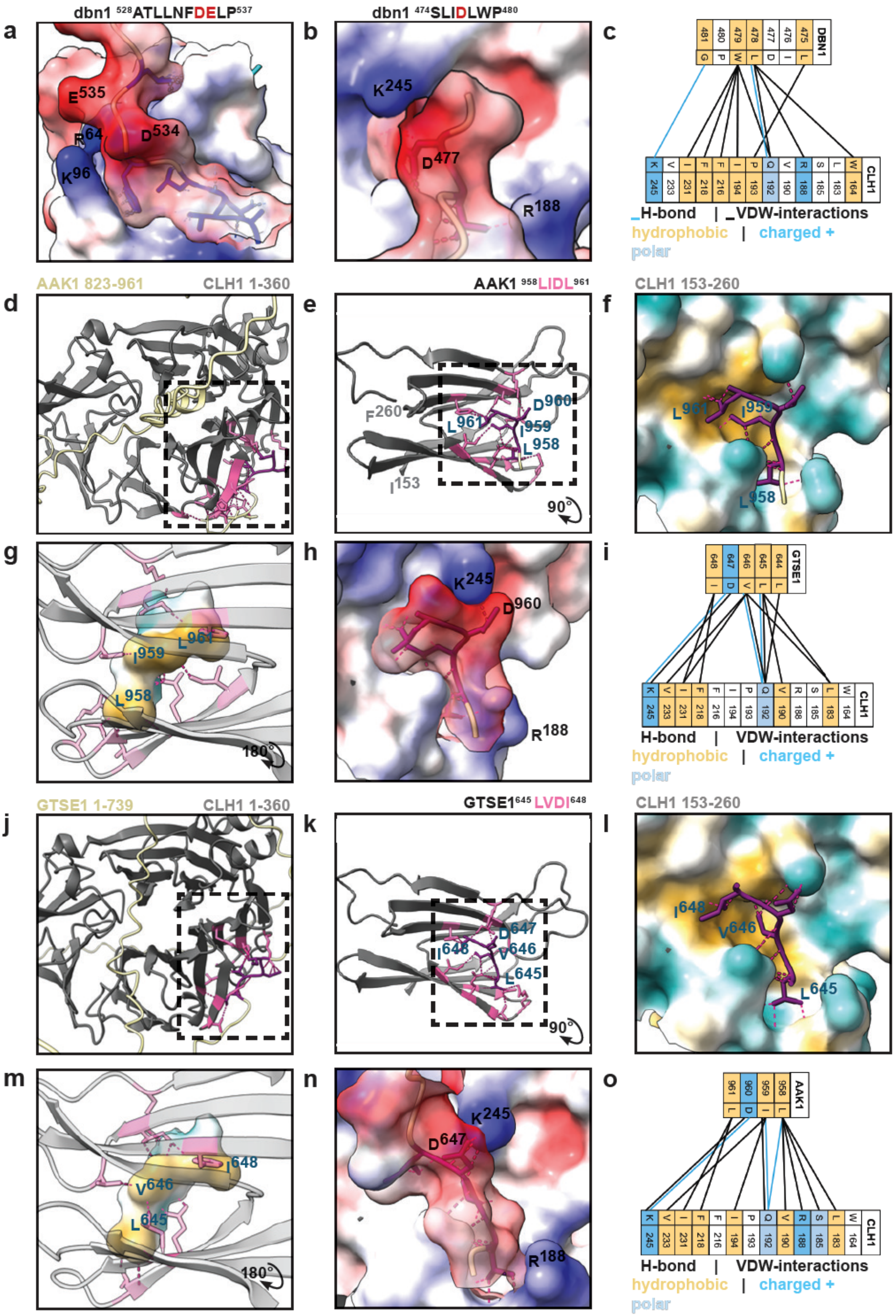

Extended Data Fig. 7

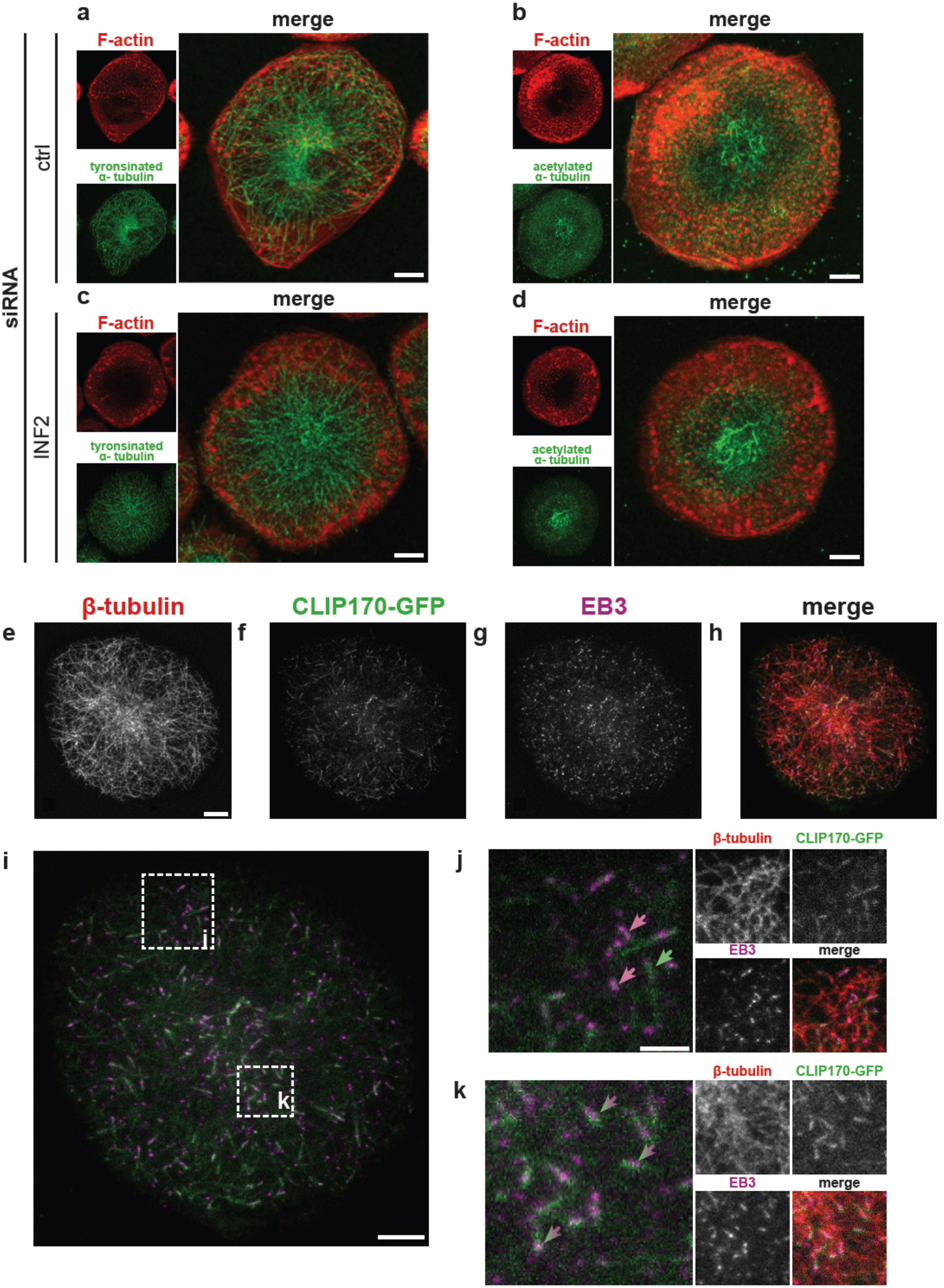

**Extended Data Fig. 8**

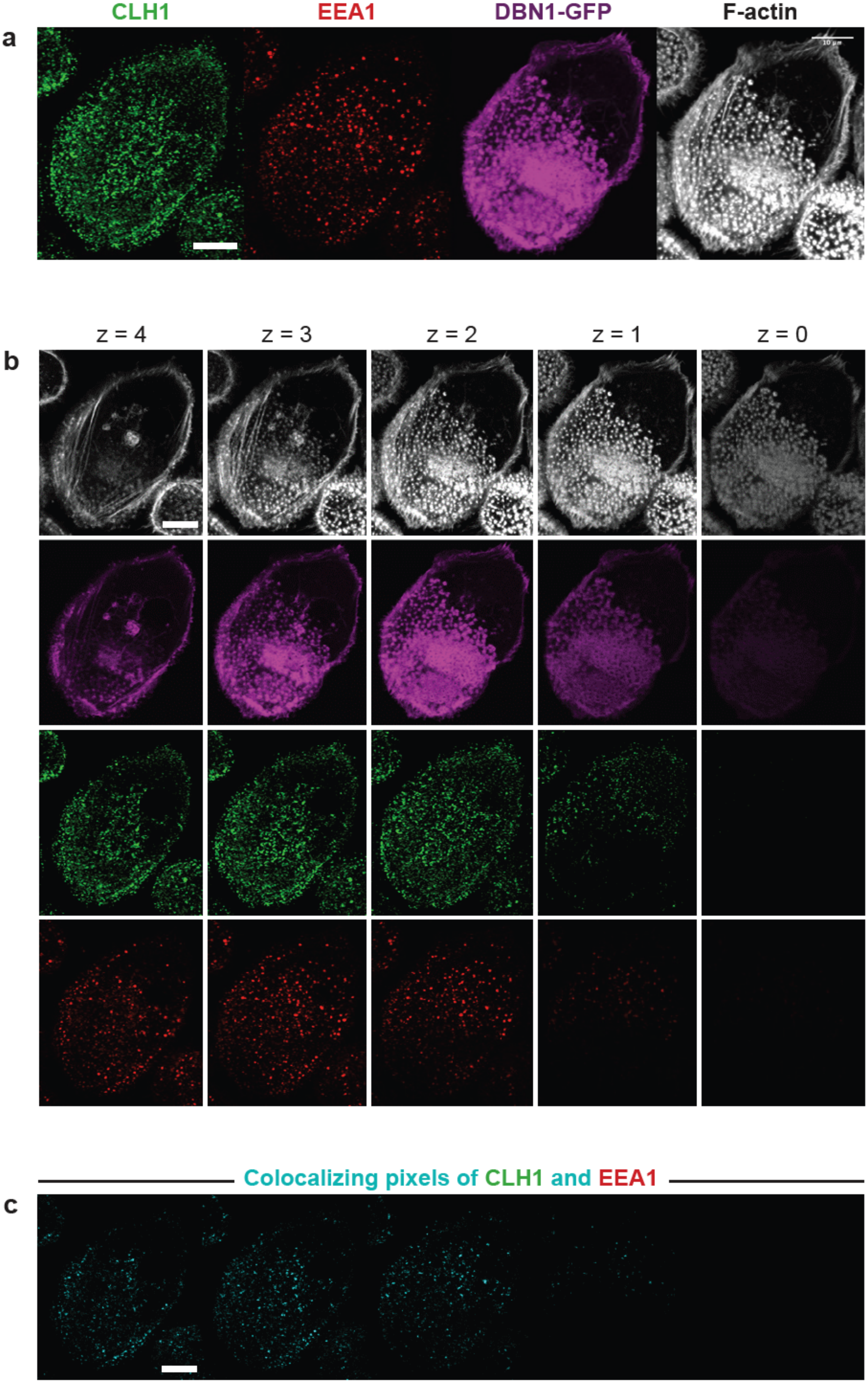

**Extended Data Fig. 9**

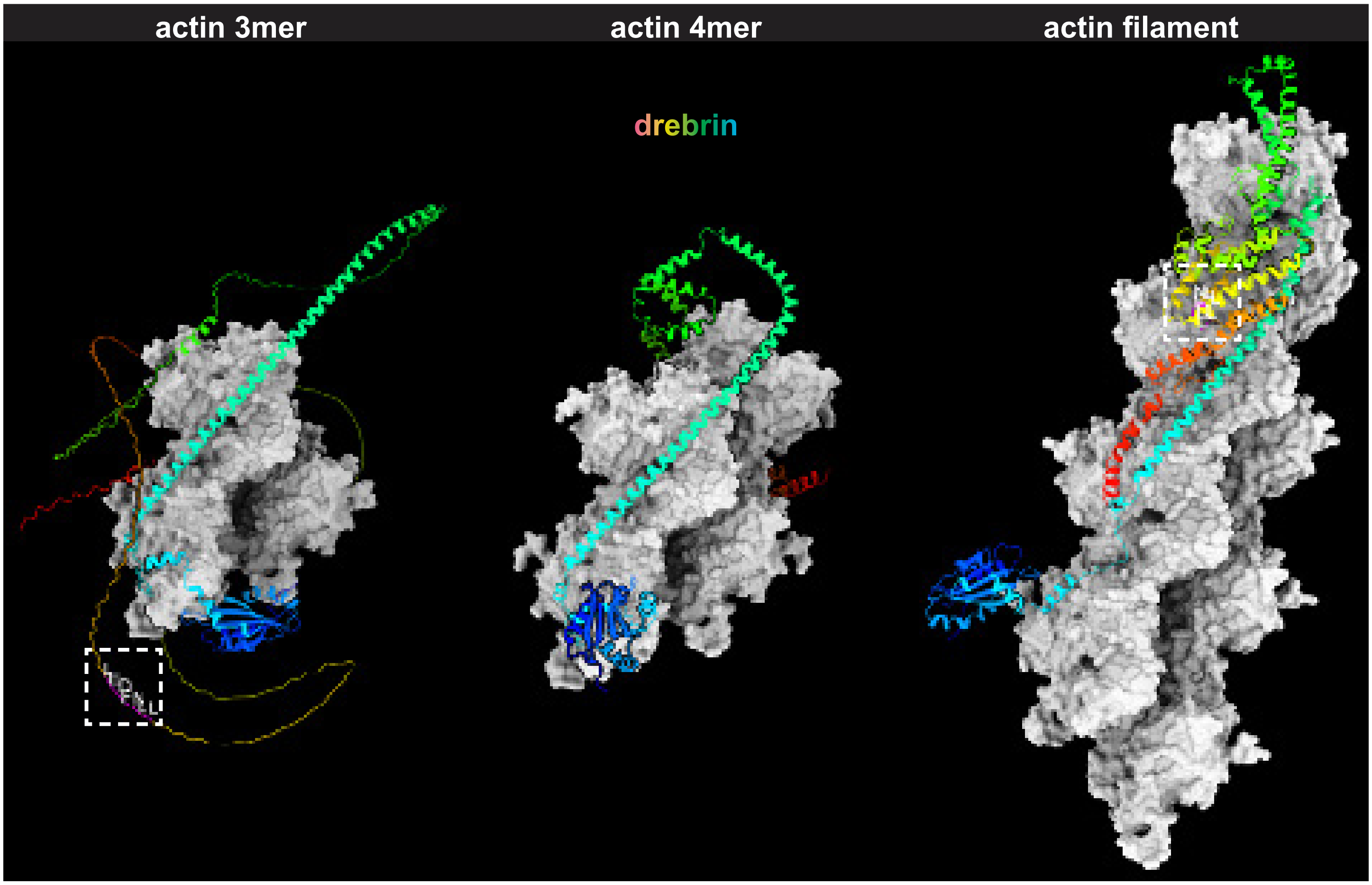
